# Size-dependence of food intake and mortality interact with temperature and seasonality to drive diversity in fish life histories

**DOI:** 10.1101/2022.08.20.504655

**Authors:** Holly K. Kindsvater, Maria-José Juan-Jordá, Nicholas K. Dulvy, Cat Horswill, Jason Matthiopoulos, Marc Mangel

## Abstract

Understanding how growth and reproduction will adapt to changing environmental conditions is a fundamental question in evolutionary ecology, but predicting the responses of specific taxa is challenging. Analyses of the physiological effects of climate change upon life history evolution rarely consider alternative hypothesized mechanisms, such as size-dependent foraging and the risk of predation, simultaneously shaping optimal growth patterns. To test for interactions between these mechanisms, we embedded a state-dependent energetic model in an ecosystem size-spectrum to ask whether prey availability (foraging) and risk of predation experienced by individual fish can explain observed diversity in life histories of fishes. We found that asymptotic growth emerged from size-based foraging and reproductive and mortality patterns in the context of ecosystem food web interactions. While more productive ecosystems led to larger body sizes, the effects of temperature on metabolic costs had only small effects on size. To validate our model, we ran it for abiotic scenarios corresponding to the ecological lifestyles of three tuna species, considering environments that included seasonal variation in temperature. We successfully predicted realistic patterns of growth, reproduction, and mortality of all three tuna species. We found that individuals grew larger when environmental conditions varied seasonally and spawning was restricted to part of the year (corresponding to their migration from temperate to tropical waters). Growing larger was advantageous because foraging and spawning opportunities were seasonally constrained. This mechanism could explain the evolution of gigantism in temperate tunas. Our approach addresses variation in food availability and individual risk as well as metabolic processes and offers a promising approach to understand fish life-history responses to changing ocean conditions.

## Introduction

The body sizes and population biomasses of aquatic species are changing rapidly in response to human-induced environmental change (Oke et al. 2020), motivating the need for mechanistic models to predict future patterns of growth and reproduction (Cheung et al. 2008, Cheung et al. 2010, Blanchard et al. 2012, Fernandes et al. 2020). Beyond thermal physiology, the consequences of changes in ecosystem productivity for life history traits (growth, reproduction, and survival) will derive from changes in prey abundance, predation risk and seasonality (Daufresne et al. 2009, Audzijonyte and Richards 2018, Neubauer and Andersen 2019). Classic and recent advances predicting optimal growth and reproduction of harvested populations (Holt 1958; Beverton and Holt 1959; White et al. 2022) have been combined with size-structured community interactions to predict the evolution of life history traits in specific taxa (Shuter et al. 2016). This approach provides a promising foundation for predicting ecological and evolutionary responses to environmental and climate change. While specific taxa within an ecosystem will adapt according to contextual factors such as phylogenetic lineage, adaptive capacity, and environmental variability, mechanistic models incorporating the roles of size-dependent foraging and predation risk, seasonality, and other metabolic processes could explain adaptive variation among closely related taxa and refine our mechanistic understanding of species’ responses to changing environmental and climate conditions (Free et al. 2019). By examining the life-histories of closely related tuna species (family Scombridae) that have invaded different environmental niches, we can gain insight into how physiological adaptations to thermal conditions, productivity, and seasonality in these factors affect allocation to growth and reproduction. This can help move us beyond broad macro-ecological patterns toward a more mechanistic understanding of the drivers of existing diversity and improve predictions of species-specific responses to future change (Alvarez-Noriega et al. 2023).

In ectothermic species with indeterminate growth, including fish, food availability and survival both increase as individuals grow larger, and growth rates can be selected to be slower or faster depending on the size-dependence of per-capita resource availability and predation risk (Conover and Munch 2002; Walsh and Reznick 2009, Hulthén et al 2021). At the same time, development at higher temperatures can lead to maturation at smaller body sizes, within and between ectothermic species (Kingsolver and Huey 2008). Allocation to growth and reproduction determine individual body sizes, fitness, and population demographic rates (Beverton and Holt 1959, Gadgil and Bossert 1970, Kozlowski 1992, Kozlowski 1996). Yet, existing theory struggles to predict how size-dependent changes in prey availability and decreases in predation risk, along with interacting effects of metabolic costs and seasonality in food availability and temperature, affect the evolution of growth (Varpe 2017). There is longstanding interest in understanding the mechanisms leading to biphasic or asymptotic growth patterns (Quince et al. 2008). The earliest growth models, including the von Bertalanffy growth model, were based on hypothesized differences in the allometric scaling of anabolism (resources taken in) and catabolism (resources spent) (von Bertalanffy 1960, Audzijonyte et al. 2022). Such growth models are routinely fit to data on size-at-age of fishes, but their mechanistic justification is necessarily simplistic; such models do not explain spatial and temporal trends in body size within and among species of fish (Audzijonyte et al. 2020), suggesting that they do not sufficiently capture the relevant intrinsic physiological and extrinsic ecological drivers of fish life histories.

The role of metabolic requirements and the trade-off between growth and reproduction in shaping growth trajectories has been given increasing attention in recent decades (Jørgensen et al. 2016, Wong et al. 2021, White et al 2022). Physiological processes can vary with the environmental temperature experienced by organisms (Brown et al, 2004, Clarke and Johnston, 1999). The allometric scaling of metabolic costs in different temperature regimes (known as the Metabolic Theory of Ecology [MTE]; Gillooly et al. 2001) has been used to predict individual body size according to the benefits of growing large (to increase foraging success and avoid predation), balanced against the costs of increased metabolic overhead and diverting resources from reproduction to growth (Wong et al. 2021, Thunell et al. 2023). More detailed dynamic models of energetic allocation have linked variable environmental conditions to growth, reproduction, and longevity (Kozłowski and Teriokhin 1999, Cichoń and Kozłowski, 2000, Lika and Kooijman 2003, Jorgensen and Fiksen 2006, Audzijonyte et al. 2022). Understandably, the results of these energetic budget models depend on specific assumptions regarding mass-based foraging success and risk of predation. Generalizing these prior results requires a unified framework incorporating metabolic demands, access to resources, and predation risk in environments of differing productivity and seasonality.

We draw on ecosystem size-spectra theory to reduce the number of *ad hoc* assumptions required about the scaling of ecological processes that arise from community interactions. We hypothesize this approach could yield more realistic predictions of diversity in fish life histories, providing significant insights into the mechanisms that determine adaptive responses to changing environmental conditions. In aquatic ecosystems, size spectra are an emergent property of community interactions (Sheldon et al. 1977, Thygesen et al. 2005, Law et al. 2009, Sprules and Barth 2016, Christensen and Andersen 2011, Andersen et al. 2016, Andersen 2019). In a community size spectrum, flow of energy between trophic levels via consumption and predation rates are characterized by individual mass, instead of species’ identity (Benoît and Rochet 2004, Blanchard et al. 2009, Andersen 2019). The key property of size spectra is that an individual’s relative position (mass) determines both its prey field (the area under the size spectrum to the left of an individual’s mass) and its risk of predation (the area to the right) (Fig. 1). That is, the size spectrum provides a simplified quantification of predator-prey interactions for a given individual in an ecosystem. Individuals are born small and grow through the size spectrum over their lifetime, consuming prey that are a fraction of their own size. At the same time, the field of predators that are capable of consuming an individual decrease as the individual increases relative to number of predators capable of consuming it, because size-spectrum theory assumes predators cannot consume prey exceeding their maximum gape. By contrast, the lower limit of prey size preference is assumed to depend on the profitability of the prey. These interactions apply to interactions between species as well as dynamics within size-structured populations of the same species (i.e., species of fish often cannibalize smaller conspecifics). Therefore, predation and consumption rates determined by different areas under a size-spectrum curve can be used to simultaneously characterize the mass-specific caloric resource availability and risk of predation experienced by an individual as it grows (Benoît and Rochet 2004, Giacomini et al. 2013, Shuter et al. 2016, Andersen 2019).

**Fig. 1.**
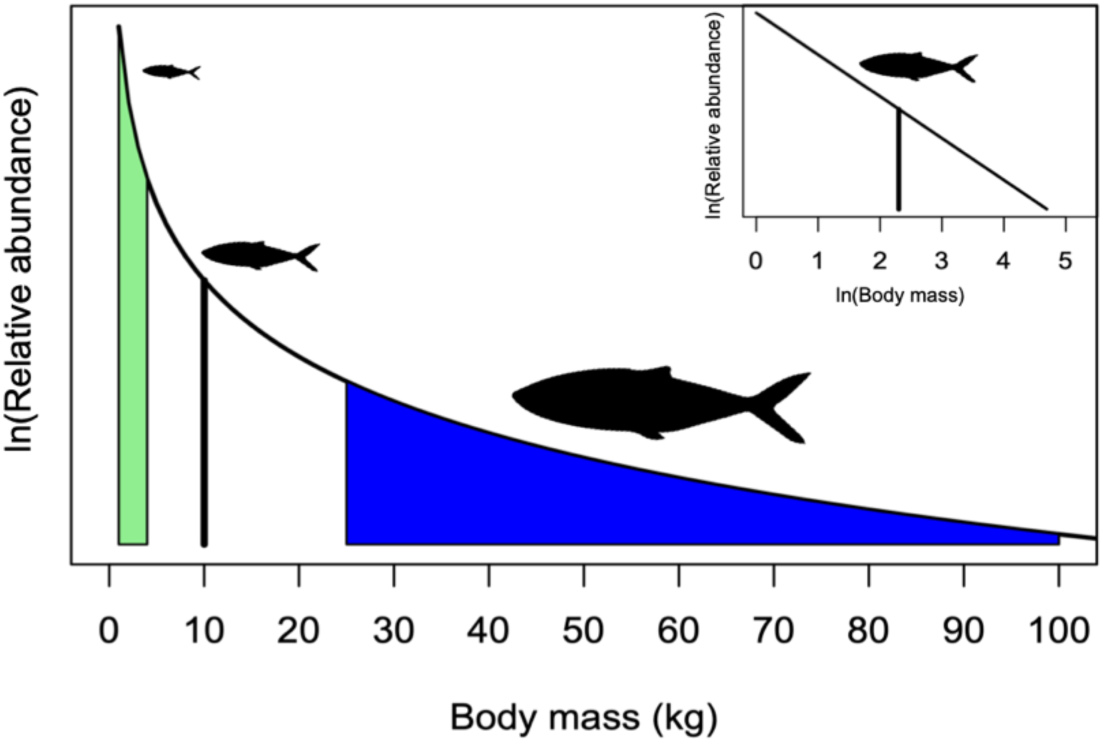
Conceptual version of the log-linear size spectrum where the prey field of an individual of 10 kg (vertical black line) is in green and the predator field is in blue. 154 Inset shows the same size spectrum in log-log space.

Our first objective was to develop an energetic model of optimal allocation to growth and reproduction that accounts for size-dependent resources and risk of predation (both derived from size-spectra) and size- and temperature-dependent metabolic requirements. We hypothesized that such a model can predict the evolution of diverse fish life histories in different environments, based on these allometric relationships. Fitness was defined as the expected lifetime reproductive output. Optimal life history strategies (allocation to maintenance, growth and reproduction) were then determined by maximizing fitness using stochastic dynamic programming (Clark and Mangel, 2000, Houston et al.1988, Mangel, 2015). We used this approach to predict the life history evolution of fishes under different scenarios of ecosystem productivity, which affects the size spectrum, and temperature.

Our next objective was to determine if our modeling approach could predict the evolution of growth, body sizes and reproductive patterns as observed in tuna species under several environmental scenarios with varying temperatures, ecosystem productivities and seasonally varying conditions. Tunas (family Scombridae) exhibit a wide range of maximum body sizes (∼ 40 cm - 400 cm), longevities (∼ 4 - 41 years) and reproductive patterns (Juan-Jorda et al. 2013, Horswill et al. 2019). Tunas are epipelagic species found in temperate and tropical waters around the world’s oceans with varying vertical, latitudinal and seasonal distributions and movements.

Confronting the predictions of our general model with data on the life history diversity observed in tuna species in different environmental conditions provided insight into the mechanisms underlying fish life histories. Where possible, our models were parameterized with values for physiological processes measured for tunas, but we also considered the sensitivity of our results to these parameters to ensure their generality across the diverse life histories observed in fishes.

## Methods

We used size-spectrum theory to infer mass-specific rates of prey encounter and mass-specific rates of predation. The relationship between the numbers of organisms *N* and individual mass *w* is a power function with a scaling parameter 𝜅 and an exponent 𝜆. This exponent has been empirically estimated to be close to -1, such that abundance is inversely proportional to mass (Sheldon et al. 1972, Trebilco et al. 2013, Sprules and Barth 2016, Hatton et al. 2021)

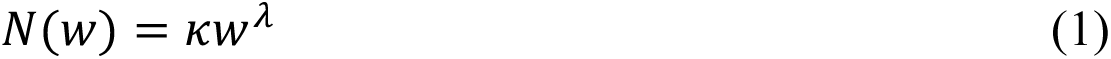

The Sheldon size spectrum, which follows from Eq. 1, is the distribution of total ecosystem biomass *B(w)* across body size classes, and is represented by 𝐵(𝑤) = 𝑁(𝑤)𝑤. Based on evidence from multiple ecosystems (Sprules and Barth 2016), the spectrum – as originally conceived (Sheldon et al 1972) and recently confirmed (Hatton et al. 2021) – is nearly flat, or may very slowly decrease as body size increases. Since 𝑁(1) = 𝜅, we view 𝜅 as a metric of ecosystem productivity, which in our model we assume is in units of kilograms per month. In a log-log plot of Eq. 1, the intercept is log(𝜅).

This phenomenon of linear aquatic size spectra emerges from three size-dependent processes: (1) the encounter rate of predators and prey; (2) the preference of predators for prey of a given size; and (3) the limit to prey consumption imposed by the size of the predator’s stomach (the predator-prey mass ratio; Benoît and Rochet 2004, Blanchard et al. 2017, Andersen 2019).

Here, we follow the modeling work of Anderson (2019), which derived a method to calculate the prey available to individuals of mass *w* by relating the productivity of a spectrum to empirical estimates of prey encounter rates, predator-prey mass ratios, and prey preferences. Andersen (2019) assumed that the prey biomass available to an individual is a concave function of mass *w*, and found it scaled to be approximately three times the size spectrum productivity 𝜅. The empirical estimates of physiological processes used to simplify this derivation can vary among species within a size class (Andersen 2019). Therefore, we assume that prey availability and individual consumption are proportional, such that the per-period food available for an individual of mass *w* is represented by

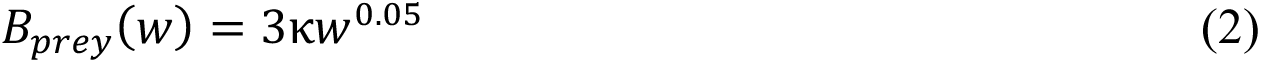

This equation approximates the integral over the range of prey biomass (in kg) available to an individual of mass *w* each month (Fig. 2A), and includes the threefold scaling factor from empirical analyses (Andersen 2019). The prey field is therefore the total biomass available to the predator, and the range of sizes of prey that it takes. This allometry is very different from the functional forms assumed in the von Bertalanffy growth model or Dynamic Energy Budget Theory. To address variability across ecosystems in productivity, we chose to vary κ in subsequent analyses, representing potential differences among taxa or other ecosystem processes.

**Figure 2.**
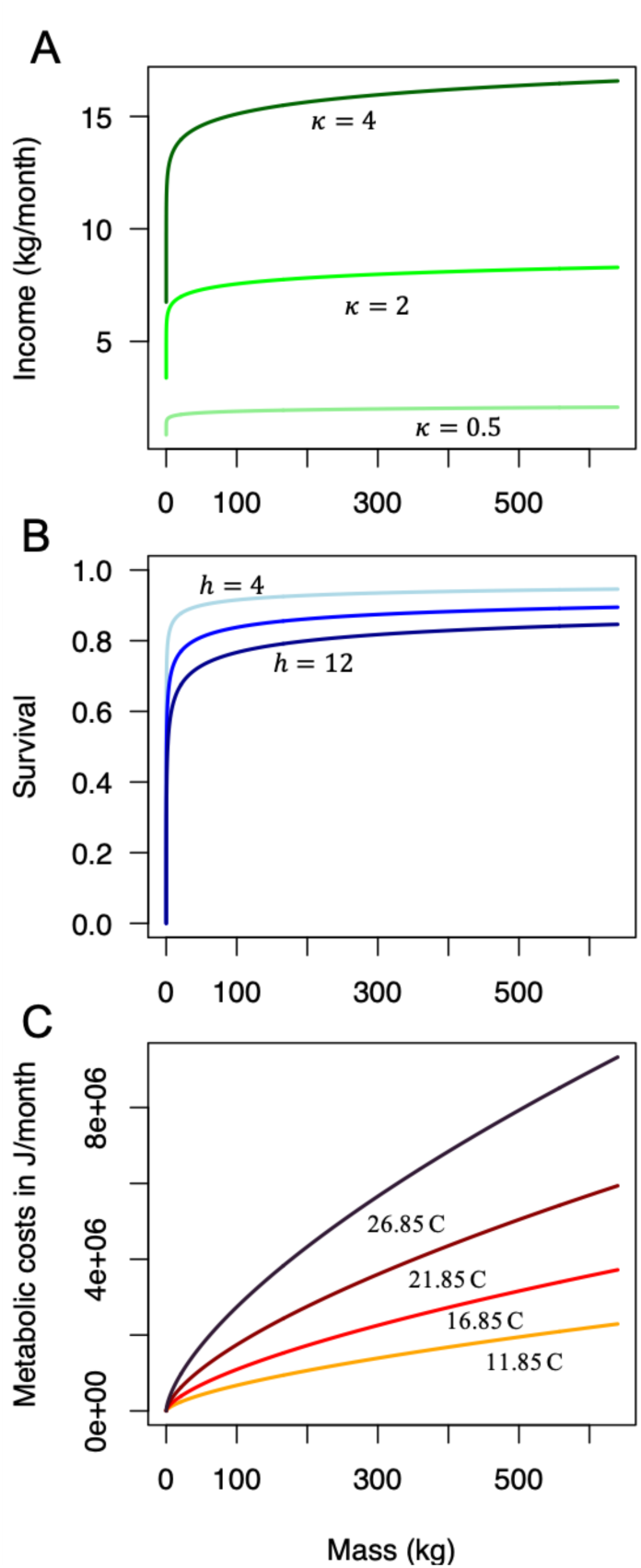
Examples of allometric relationships between individual mass and (A) income or energy gained (in kg per month) from prey biomass for differing ecosystem productivities κ (Eq. 2); (B) the scaling of survival (the inverse of the size-based risk of mortality via predation) and its interaction with predator *h* (Eq. 4), such that higher values of *h* represent more efficient predators; and (C) metabolic costs (Eq. 5), which also scale with temperature 𝜏 (on the graph, it is presented in degrees Celsius, ℃; values in Kelvin are 285, 290, 295 and 300K). Note that these curves represent a subset of values considered in our results.

Second, we calculate survival per time from the mass-specific risk of predation that emerges from size-spectrum theory. We assume that the processes determining consumption rates of predators in the size spectrum can be used as a proxy for individuals’ instantaneous rate of mortality from predation, following the derivation in Andersen (2019). This derivation is based on first principles. For gape-limited taxa like fishes, a predator’s prey field depends on its encounter rate with prey in its preferred size range. This encounter rate (and the clearance rate) can be measured in units of volume per time, as aquatic species forage in a three-dimensional habitat (Kiørboe and Hirst 2014). The risk of mortality from a single predator is therefore the volume of prey cleared by a predator, relative to the volume encountered, and weighted by the size of its preferred prey. This must be multiplied by the density of predators and integrated over all sizes (Box 2, Andersen 2019). For simplicity, we ignored potential effects of temperature on encounter or clearance rates that could arise from an increase in activity associated with warmer environmental conditions. In Eq. 3 (defined below) we used an empirically estimated constant of 0.07 to characterize the scaling of prey vulnerability with its mass (Andersen 2019), which is based on estimates of preference windows of predators and the volume of water each clears per month that come from empirical distributions of prey sizes in predator guts (Ursin 1973). We also define a coefficient *h,* which modifies the probability of consumption - how likely a predator is to capture the focal individual (based on its capture efficiency) - and used the reported allometric exponent of -0.25 to represent how predation mortality scales with body size (Fig. 2B; Eq. 2.11 in Andersen 2019). Anderson (2019) integrated over the Sheldon spectrum to produce a relationship predicting the scaling of mortality risk for prey 𝜇_-_(𝑤) per month:

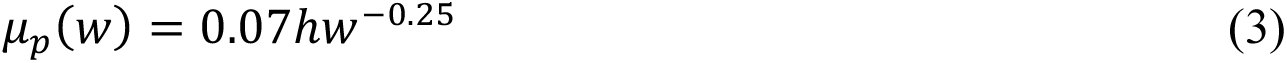

We convert this instantaneous rate to the probability of escaping from predators during each time interval (Hilborn and Mangel 1997, Mangel 2006), so we can represent the probability an individual survives 𝛾_pred_ (𝑤) as:

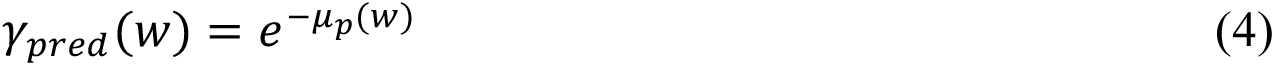

The resulting mass-specific survival through each month is plotted for different values of *h* in Fig. 2B.

### Defining metabolic costs that depend on temperature and body mass

Our energetic model includes metabolic costs which depend on temperature and body mass. We assumed that metabolic costs increase with body mass and environmental temperature (Clarke 2006). We modelled individual costs 𝐶(𝑤, 𝜏) (in joules) as a function of temperature 𝜏 in Kelvin, following the general form introduced in the MTE (Gillooly et al. 2001):

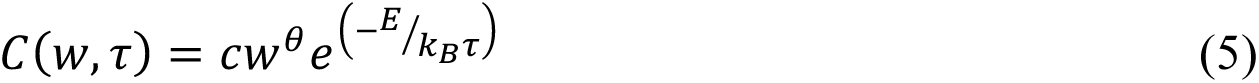

Evidence for the MTE suggests the activation energy *E* (the energy required for the reactions of respiration and other metabolic processes) varies little among taxa (Brown et al. 2004, Bernhardt et al. 2018, but see Lindmark et al. 2022); Boltzmann’s constant 𝑘_N_ also does not vary. The normalization coefficient *c* accounts for differences among taxonomic groups in the intercepts of the linear relationship that arise from second-order effects such as stoichiometry or respiratory surface areas (Bigman et al 2021). The slope of this relationship in log space, *θ*, is strikingly similar among taxa (Brown et al. 2004). We used a value for *θ* estimated from physiological studies on tunas (Table 1) but varied it in sensitivity analyses. Note that 𝜏 in Eq. 5 is traditionally in Kelvin. However, we rescale all values in our results in units of Celsius. We use this general formula for scaling of metabolic costs at different temperatures 𝜏 to describe the monthly energetic expenses of an individual of mass *w* (Fig 2C). Defining the relationships in Eqs. 2, 4, and 5 allowed us to specify mass-dependent survival and energy dynamics and therefore examine the variables influencing growth and reproduction in a common currency of individual fitness. All parameters are defined and their values reported in Table 1.

**Table 1.** Description of parameters and variables.

| Parameter or Variable | Description | Value |
| --- | --- | --- |
| $w$ | Body mass in kg | - |
| $N$ | Number of individuals in a size category (or trophic level) when considering a Sheldon size spectrum | - |
| $B$ | Absolute biomass in a size category (or trophic level) when considering a Sheldon size spectrum | - |
| $\kappa$ | The spectrum parameter, which defines the total biomass of organisms of the smallest body size $w$ in a given ecosystem; Andersen (2019) gives an estimate of 10 gained by averaging over all Predator-Prey Mass Ratio estimates measured from gut contents. We vary it to represent ecosystem differences in overall ecosystem productivity | Varies |
| $B_{prey}$ | Biomass of prey expected by a focal individual | - |
| $\mu_p$ | Instantaneous risk of mortality due to predation, which depends on body mass and position in the size spectrum | - |
| $h$ | Predation risk, comprised of predator satiation estimates (estimated from gut contents) and predator preference (or effectiveness) for consuming prey of a given mass (Andersen 2019) | 4, 8 |
| $\tau$ | Temperature of the environment (in degrees Kelvin except where noted) | 285-300 |
| $C$ | Metabolic requirements (costs) that scale with mass and temperature | - |
| $c$ | Normalization constant scaling metabolic costs (in J), based on metabolic rate data from tunas (Kitchell et al. 1978) | $5 \times 10^{16}$ |
| $k_B$ | Boltzmann constant, relating particle energy to temperature in units of $m^2 kg s^{-2} K^{-1}$ | $1.3 \times 10^{-23}$ |
| $E$ | The average activation energy for the rate limiting enzymes in metabolism in units of joules; from the metabolic theory of ecology (Brown et al. 2004). | $1.04 \times 10^{-19}$ |
| $\theta$ | Metabolic scaling exponent; values vary among clade, here we use a value reported for tunas (Clarke and Johnston 1999) | 0.66 |
| $\rho$ | The energy density of tuna body mass in our model in J/kg (estimated empirically and reported in Chapman et al. 2011) | $4.2 \times 10^6$ |
| $t$ | Time in monthly time steps in the dynamic model | - |
| $T_{max}$ | Maximum lifespan in months | 216 |
| $l, L(t)$ | Body length (in cm) This is a dynamic state variable but can only increase with time. The maximum value possible is 400 cm. For a specific value, we use the capital letter notation. | - |
| $s, S(t)$ | Lipid stores (in joules) – this is a dynamic state variable representing energy stores that can be used for metabolism, growth, and reproduction. | - |
| $a$ | Scale coefficient relating length to structural mass, similar to values estimated empirically for bluefin tuna and reported in Pignalosa et al. (2020). | $1.0 \times 10^{-5}$ |
| $w(t)$ | Structural mass of the individual (in kg) at time $t$ – this depends on $L(t)$ | - |
| $v$ | The fraction of structural mass that determines the critical threshold of energetic mass needed to avoid starvation | 0.1 |
| $\varphi$ | The fraction of structural mass that determines the maximum possible reproductive output in a monthly time step | 0.2 |
| $\gamma_{pred}$ | Survival from predation from one month to the next, emerges from risk of mortality | |
| $\gamma_s$ | Survival, given that the individual has sufficient energy reserves to meet metabolic requirements and avoid starvation | |
| $g$ | Proportion of lipid stores allocated to growth (this allocation decision is optimized by the dynamic programming equation); can take values between 0 and 1 | |
| $r$ | Proportion of lipid stores allocated to reproduction (this allocation decision is optimized by the dynamic programming equation); can take values between 0 and 1 and the sum of $g$ and $r$ cannot exceed one. | |
| $R(l, s, t)$ | Current fitness, the product of lipid stores and the optimal allocation to reproduction, $r$ | |
| $V(l, s, t)$ | Expected accumulated lifetime reproduction from time $t$ onwards, given that $L(t) = l$ and $S(t) = s$ | |

We note that environmental temperatures can also affect the encounter rate of prey through its effects on activity level. Additionally, the productivity of ecosystems is expected to increase with temperature as it increases rates of consumption and respiration, improving growth rates. We have chosen to address these potential interactions between temperature and energetic intake by comparing growth patterns that occur at the same temperatures with differing levels of ecosystem productivities, such that the potential relationship between 𝜅 and temperature is implicit.

### Defining fitness and determining the optimal allocation strategy

To find the optimal allocation strategy in different scenarios of ecosystem productivity and temperature conditions, including seasonal variation in the scenarios, we developed a model of an individual’s energy budget, tracking two physiological state variables, the body length *l* and energy (lipid) stores *s* of an individual, which vary dynamically over an individual’s lifetime (Fig. 3). Following the conventions for dynamic state-variable models, we denoted the state variables *l* and *s* as lowercase in the dynamic programming equation, representing the fact they are iterated values; potential future states are denoted as *l’* and *s’*. Later, when we refer to values of the state variables at a specific time, we use uppercase *L* and *S*.

**Figure 3.**
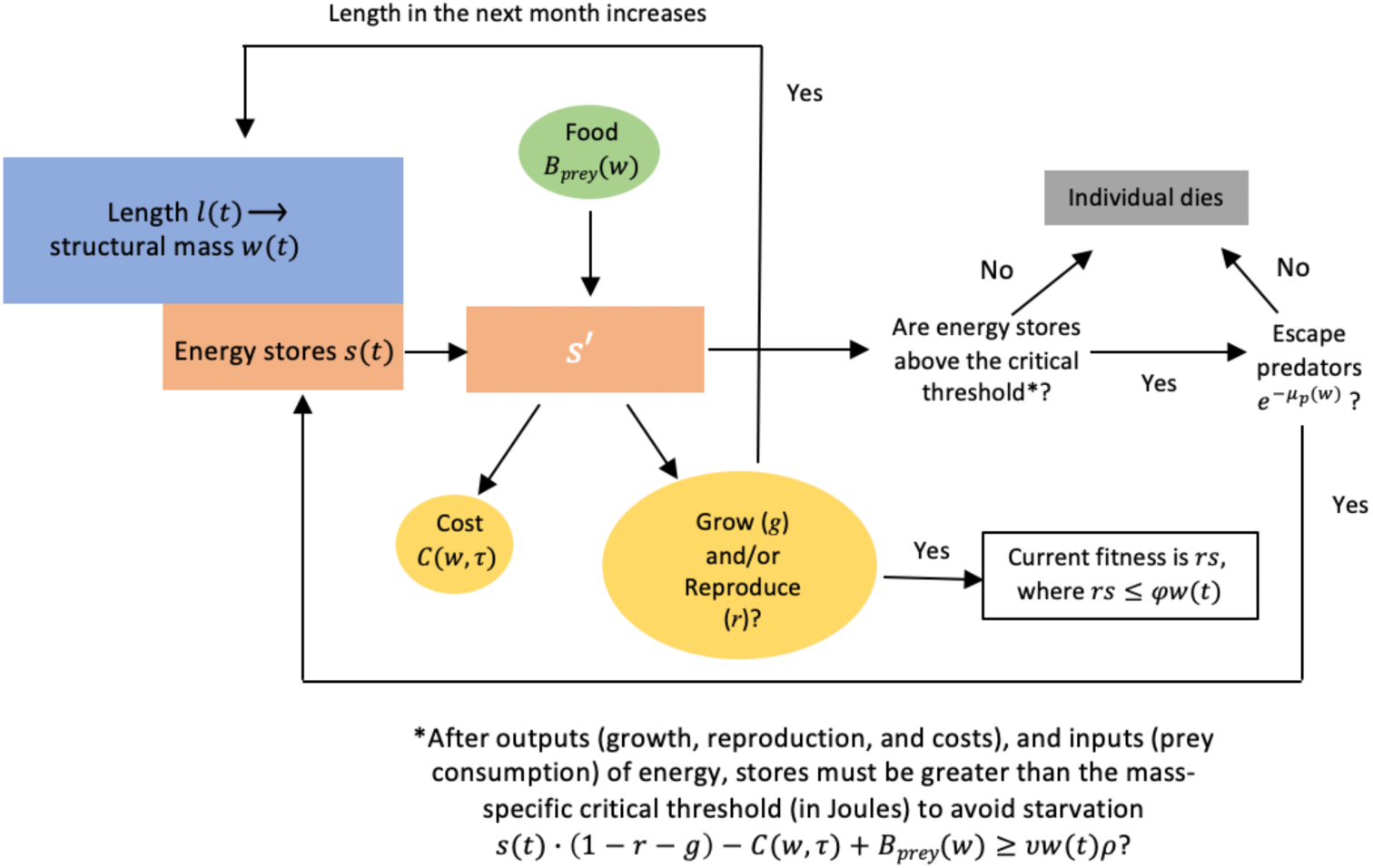
Conceptual overview of the optimization algorithm calculating the two dynamically varying state variables, body length 𝑙(𝑡) and energy stores 𝑠(𝑡) within each month of an individual’s life *t,* as well as current and expected fitness. Arrows represent the flow of energy or decisions. Round shapes represent energetic input (green) and outputs (yellow); rectangular shapes are model states and outputs (fitness and fate). Note that 𝑙(𝑡) and 𝑤(𝑡) are related through Eq. 6. Both 𝐵_𝑝𝑟𝑒𝑦_ and 𝜇_-_ are determined by the size spectrum.

We define fitness as expected lifetime reproductive output, averaged over the stochastic process of mortality from both predation and starvation, which we calculated numerically using stochastic dynamic programming (Houston et al. 1988; Clark and Mangel 2000, Mangel, 2015). This method allowed us to consider how individual age and physiological state (energy stores and body length) affect the optimal trade-off between growth and reproduction in the context of expected lifetime reproductive output. We assumed that an individual allocates a proportion of its budget to growth and reproduction on monthly time intervals. Our choice to model allocation to growth and reproduction as proportions of an individual’s energy budget builds on prior dynamic state-variable models of fish growth (Jørgensen and Fiksen 2006, Chapman et al. 2011).

Given specific values of 𝐿(𝑡) and 𝑆(𝑡), representing individual length and energy stores at the start of month *t,* structural mass 𝑊(𝑡) is related to length with a standard cubic function (Froese 2006):

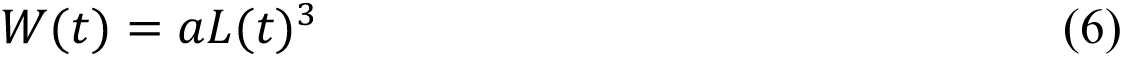

We assumed that only structural mass is relevant for size-dependent gains (𝐵_𝑝𝑟𝑒𝑦_) and costs (*C*) (Fig. 3). We convert from mass to units of energy (joules) and back using a conversion factor estimated for tunas (𝜌 = 4.2 × 10^_^J/kg; Chapman et al 2011) but that could take other values for other species. Each month, an individual acquires energy from food, determined by their structural mass *w* via Eq. 2. We then allowed the individual to allocate proportions of its energy budget (which includes income and any stored energy) to growth (*g*) and reproduction (*r*).

Given an individual with specific values for length 𝐿(𝑡) and energy stores 𝑆(𝑡), we calculated the increment of growth expected for each possible proportional allocation *g* by converting the fraction of lipid stores from joules to the equivalent mass 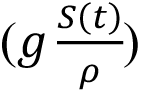, and then adding it to existing mass, such that 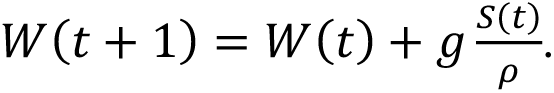. We then calculated the new length by rearranging the mass-length relationship 𝑊(𝑡) = 𝑎𝐿(𝑡)^3^:

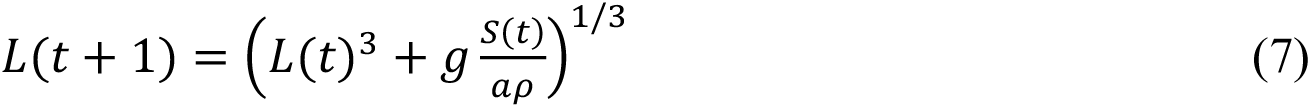

We repeated this for every possible combination of values of the state variables, *l* and *s.* Values for proportional allocation of lipid stores to reproduction *r,* along with *g,* combine to determine the dynamics of energy stores and length from one month to the next:

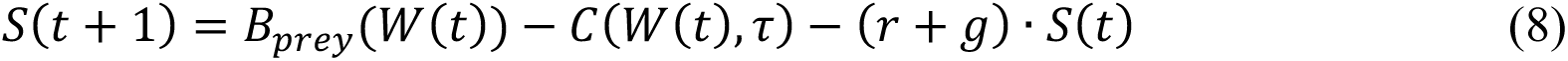

We assumed that both stored energy and reproduction were limited by an individual’s structural mass (which in turn depends on its length). These constraints represent limits on both the amount of lipid can be stored, and the mass of gametes that can be produced, given the capacity of the body cavity. If the proportions of energy allocated to reproduction and growth were less than 100% (meaning 𝑟 + 𝑔 < 1), the remaining energy was assumed to be stored for future use, given that total reserves did not exceed 60% of structural mass. This value was arbitrary, but exploratory analyses suggest it did not have a strong effect on the results presented here because in practice individuals do not store their energy long enough to come close to exceeding this limit. Reproductive output (in units of kg) was similarly constrained so that it cannot exceed a fixed proportion of 𝜑 of structural mass 𝑤(𝑡), such that the following condition must be met:

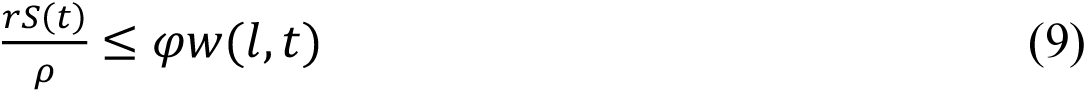

This size-based limit on total reproduction is used in the calculation of current fitness.

### Expected future fitness

At any age, expected fitness was the sum of current reproduction and accumulated future reproduction, which was calculated assuming the individual behaved optimally for all future ages. This required calculation of the future states (length and lipid stores) given each combination of allocation to growth and reproduction. We denoted potential values of future states as 𝑙^m^and 𝑠^m^. These are

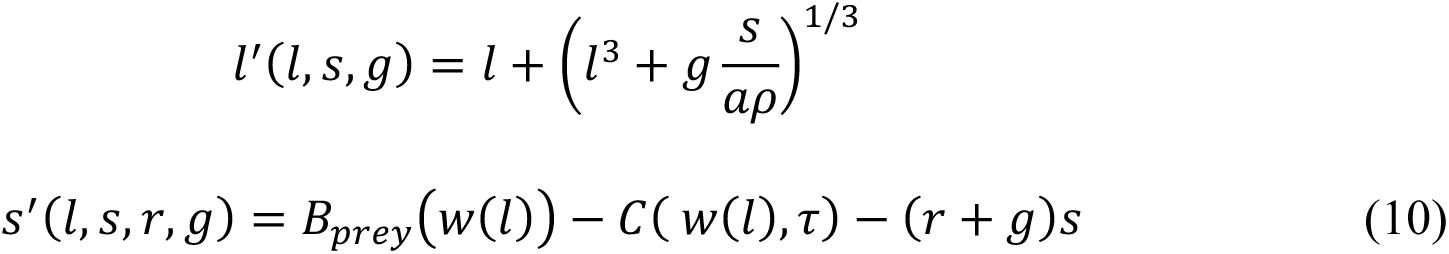

where 𝑤(𝑙) is the structural mass of an individual of length *l.* At the end of the month, if an individual’s lipid stores fell below the critical threshold for its mass (its expenditures have exceeded its energy budget), it starved (Suppl. Fig. 2). We let 𝛾_r_(𝑠, 𝑙) denote the probability of avoiding starvation in the current time and modelled it as

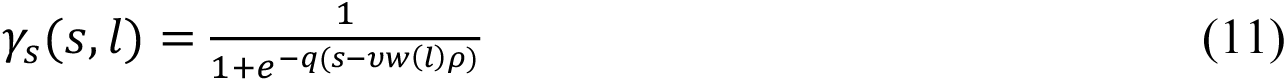

involving two new parameters 𝜐 and *q.* The parameter 𝜐 determined the level of stores at which starvation begins; *q* is a shape parameter. When 𝑠 = 𝜐𝑤(𝑙)𝜌, the right-hand side of Eq. 11 is always ½. When *q* is large, then the right-hand side of Eq. 11 is approximately one when 𝑠 > 𝜐𝑤(𝑙)𝜌 and zero otherwise. Thus, this function specified that if combined allocation to growth and reproduction caused an individual’s lipid stores to decrease below the critical value for its mass, its probability of survival decreased smoothly toward zero (See Suppl. Fig. 1 for more details).

We varied growth *g* and reproduction *r* and determined the combination that maximized fitness. Equations 7-11 define changes in individual state and in fitness for all allocation strategies (all values *r* and *g*). With these functions in place, we can find the allocation strategy that maximizes current and future fitness at every age until the age of senescence or maximum lifespan, *T*, is reached (for all scenarios, we assumed the maximum lifespan of *T =* 216 months or 18 years).

We define 𝑉(𝑙, 𝑠, 𝑡) as the maximum expected accumulated reproduction between time *t* and *T,* given size 𝐿(𝑡) = 𝑙 and lipid stores 𝑆(𝑡) = 𝑠. Since there can be no accumulated reproduction after *T,* we assumed 𝑉(𝑙, 𝑠, 𝑇) = 0. Expected future fitness at every age *t* < *T* was found by solving the stochastic dynamic programming equation, which for all values of allocations *r* and *g*, and age in months *t* decomposes expected reproduction from time *t* onwards into reproduction at time *t* and expected reproduction from time 𝑡 + 1 onwards given the new values of the states:

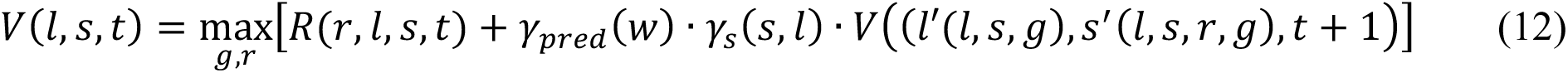

The first term of the on the right-hand side of Eq. 12 represents reproduction in month *t*. The second term represents expected future reproduction, discounted by the probability of surviving predation 𝛾_pred_ (𝑤) and starvation 𝛾_s_(𝑠, 𝑙), When these are combined, we can obtain expected lifetime reproduction from time *t* onwards, given that size 𝐿(𝑡) = 𝑙 and lipid stores 𝑆(𝑡) = 𝑠. The dynamic programming algorithm (Mangel and Clark 1988; Houston and MacNamara 1999) iterates over all viable combinations of *l* and *s,* at each time *t,* and stores the fitness of each allocation strategy. The optimal strategy (marked with an asterisk) at time *t* is the combination *g\** and *r\** associated with the greatest current and future fitness. Further details of the optimization algorithm are given in the Appendix. In Suppl. Fig 3 we illustrate the array for both allocation strategies (*g*(l,s,t)* and *r*(l,s,t)*) at two ages, for all possible combinations of length and lipid stores.

### Calculating the fates of a cohort of individuals allocating optimally

Assuming that an individual followed optimal allocations determined by Eq. 12, we specified the length at birth and used forward iteration (Clark and Mangel 2000) to determine the accumulated mortality and reproductive output as a function of time. Some combinations of states (length, lipid stores and age) will not arise naturally and others are inviable (the dark blue regions of Suppl. Fig. 3). We recorded body lengths and reproductive outputs in subsequent month, and calculated the probability of survival to age, given both the risk of starvation and the risk of predation. We defined expected lifespan as the age past which expected survival was less than 3%. For simplicity, we considered reproductive output in units of energy (joules) rather than considering allocation to offspring size and number (Kindsvater et al. 2010); this can include migration costs. We did not build in any assumptions about age or size at maturation, but rather let maturation patterns, along with natural trade-offs between growth and reproduction (Suppl. Fig. 3) emerge from patterns of allocation.

### Scenarios for environmental variation

We used our general energetic model to ask whether we could predict a range of fish life histories (patterns of growth, reproduction, and lifespan) that are evolutionarily advantageous across different scenarios of ecosystem productivity and environmental temperatures. To do this, we developed different productivity and temperature scenarios corresponding to different conditions that impose different metabolic costs to individuals according to Eq. 5. We solved for the optimal life histories under different environmental temperatures (𝜏 in Eq. 5 in units of Kelvin), converted to Celsius and ranging from 11.85 to 26.85 ℃ (285-300 K) in five-degree intervals. Temperature affected energetic budgets of individuals, but did not directly affect consumption rates, which increase in warmer conditions (Clarke 2006). To tease apart the effects of thermal costs from temperature-effects on resources, we considered these temperature ranges in different scenarios for ecosystem productivity, for which 𝜅 ranged from 0.25 to five, in factorial combinations.

### Case study predicting life history diversity of tunas (Scombridae)

We asked if our general energetic model predicted the patterns of life history variation observed in tunas, a group of species adapted to different environments. There are 15 species of tunas within the Family Scombridae, from five genera: *Allothunnus*, *Auxis*, *Euthynnus*, *Katsuwonus*, and *Thunnus* (Collette et al. 2001). These species inhabit a wide range of environmental conditions in marine ecosystems. Paleo-oceanographic evidence suggests that ancestral tunas evolved in a tropical environment approximately 60 million years ago (Monsch 2000) and over time they have diversified and evolved a suite of morphological and physiological adaptations that have allowed them to expand their distributions into more temperate environments or deeper colder waters where they can encounter higher prey densities to support their high somatic and gonadal growth rates (Dickson and Graham 2004). Currently, tunas can be found in coastal and oceanic pelagic waters, and have wide geographic distributions, ranging from the tropics to higher temperate latitudes with some degree of habitat partitioning by depth. Tropical tunas can spawn throughout the year, while the subtropical and temperate tunas undergo seasonal migrations returning form cool high latitude feeding grounds back to warm waters for spawning (Juan-Jordá et al 2013, Horswill et al 2019). Reflecting their tropical ancestor, all tunas (except for the slender tuna *Allothunnus fallai*) spawn in warm waters, requiring a sea surface temperature of at least 24 ℃ (Schaefer 2001). This is a key aspect of their reproductive biology that we include in our model scenarios.

To connect our general energetic model more explicitly with the observed patterns of life history variation in tunas, we followed a proposed categorization of tunas into three ecological lifestyles (Bernal et al. 2017). The three general lifestyles are based on species-specific vertical, latitudinal, and temporal (seasonal) distributions and movement patterns of tunas (Figure 4; Bernal et al. 2009, Bernal 2011, Bernal et al. 2017). The first ecological lifestyle represents a tuna species that largely remains within the warmer and well-oxygenated surface layer above the thermocline (generally above 20 ℃) during both day and night. These tuna species have limited vertical movements as they do not descend below the thermocline (Figure 4). Coastal species, such as the tropical frigate tuna (*Auxis thazard*) may typify this group. The second ecological lifestyle represents tuna species that spend the majority of the time above the thermocline (generally above 20 ℃) but also visit depths below the thermocline for foraging (Figure 4). The oceanic species of yellowfin tuna (*Thunnus albacares*) with year-round tropical distributions typifies this group. Its vertical movement exposes this species to a wider range of temperatures and to less-oxygenated waters at depth but only for short periods of time because this species is not hypoxia tolerant (Schaefer et al. 2009). The third ecological lifestyle characterizes tuna species that are exposed to a wider range of environmental conditions and spend significant periods of time in colder waters (Figure 4). The oceanic and temperate Atlantic bluefin tuna (*Thunnus thynnus*) are one species that typifies this group, spending most of the year at higher latitudes in colder and more productive waters between the upper mixed later and the cooler deep waters below the thermocline, and migrating to warmer waters for spawning (Bernal et al. 2017). To examine whether our general model could predict these three broad ecological lifestyles of tunas, we modelled environmental scenarios that correspond to the habitat of each representative species in terms of temperature, ecosystem productivity, and seasonality (Table 2). We chose parameters based on the results of our general model, but also added seasonal fluctuations in environmental temperatures (which increased metabolic costs) and productivity to represent the bluefin tuna migration from temperate, productive waters to their tropical spawning grounds. We then asked whether the life-history traits emerging in these scenarios are consistent with the range of reported sizes of three representative species corresponding to each lifestyle (Juan-Jordá et al. 2016). This analysis assumes that our model assumptions regarding the relationship between temperature and metabolic costs, which is derived from macro-ecological patterns (Gillooly et al. 2001), holds for closely related species of tunas. Further, note that we do not have direct information on values of 𝜅 in different oceanic environments. Variation in 𝜅 in this analysis could represent positive effects of temperature on overall resource availability as well as consumption.

**Figure 4.**
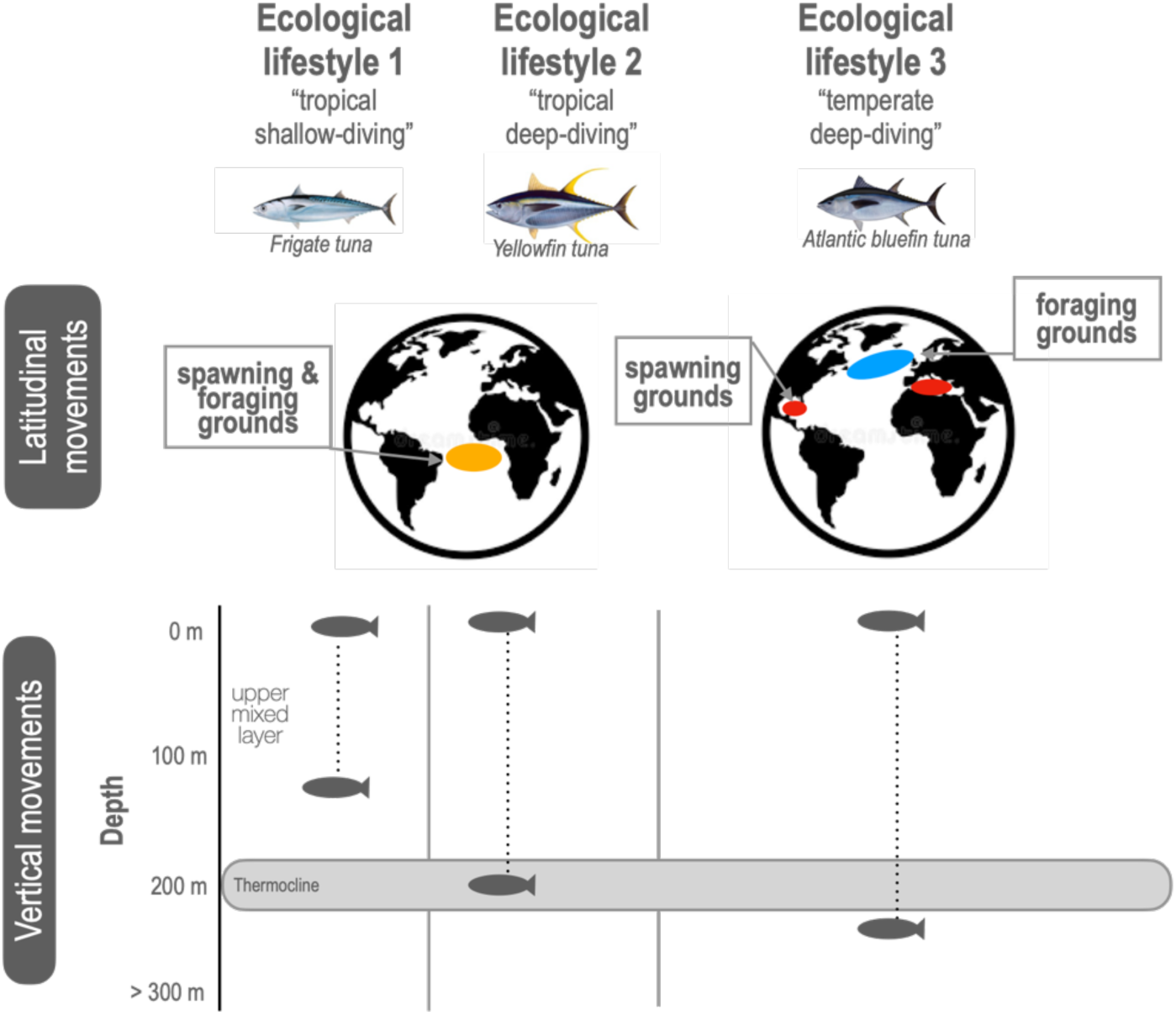
Three representative ecological lifestyles of tunas and their distribution patterns based on their latitudinal and vertical movements. Characteristic spawning and foraging grounds are shown for each lifestyle; The three ecological lifestyles illustrate a tropical shallow-diving frigate tuna *Auxis thazard*, a tropical deep-diving yellowfin tuna *Thunnus albacares,* and a temperate deep-diving Atlantic bluefin tuna *Thunnus thynnus*. Fish silhouettes represent the depth distribution where species spend most of their time. The thermocline is defined as the depth range within which the water temperature changes rapidly and separates the water column into the upper well-mixed surface layer (water above 20 ℃) and the deeper waters (waters below 15 ℃). Figure modified from Bernal et al. 2009 and Bernal et al 2017.

**Table 2.** Parameter values for environmental scenarios corresponding to the three ecological lifestyles of tunas described in Fig. 3, with corresponding predicted body size and maximum observed fork length (cm) for three representative species (data from Juan-Jordá et al. 2016).

| Lifestyle | Environment | Baseline or winter temp | Yearly mean $\kappa$ | $h$ | Predicted body size | Representative species | Reported range of body sizes |
| --- | --- | --- | --- | --- | --- | --- | --- |
| Tropical shallow | Constant | 26.85 C | 0.1 | 8 | 62 cm | Frigate tuna | 40-60 cm |
| Tropical deep diving | Constant | 21.85 C | 1 | 8 | 214 cm | Yellowfin tuna | 125-231 cm |
| Temperate deep diving | Seasonal | 11.85 | 2.5 | 12 | 361 cm | Bluefin tuna | 203-427 cm |

### Sensitivity analyses

We ran a series of tests to examine how our choices of parameters in the size spectrum influence model predictions. Specifically, we varied values of *h* (representing efficiency in prey capture) in the function describing the risk of mortality (Eq. 3) and 𝜑 (the fraction of body mass that can be devoted to reproduction) in the reproductive constraint (Eq. 9). We also conducted sensitivity analyses in which we varied the scale and shape coefficients in the metabolic cost function (*c* and 𝜃, respectively, in Eq. 5). Motivated by our tuna case study, we also varied seasonality in resources, thermal (metabolic) costs, and spawning. We considered seasonal environments in which only resources and only thermal costs varied, to understand how these factors alone contributed to observed variation in body size and reproductive output. We additionally considered growth and reproductive patterns with an extended warm season (where individuals could spawn in warm temperatures for six months of the year, instead of three). Finally, we determined in preliminary analyses that the maximum lifespan *T* did not strongly affect model results, because most individuals reach a maximum body size well before this threshold.

## Results

An asymptotic growth pattern naturally emerged from the model, after a period of exponential growth early in life (Fig. 5). Our model predicted age-specific relationships between body size (length, from which we calculate mass using Eq. 6) and reproduction that correspond to a range of fish life histories (Fig 5). In general, we found that individuals allocated energy to growth early in life and shifted this allocation to reproduction in later life. Ecosystem productivity (𝜅) alone generated a range of maximum body sizes, from less than 100 cm at low levels of productivity to well over 350 cm (Fig. 5A). Across scenarios of ecosystem productivity, the predicted trajectories for individual growth were identical in early life (before age 2); subsequent growth slowed earlier in lower productivity environments (Fig. 5A). For all life histories, allocation to reproduction began at very low levels sometime during the individual’s second year (Fig. 5B) and increased steadily as the individual aged. Both the rate at which reproduction increased with size and the maximum reproductive output were consistently greater with increased ecosystem productivity (Fig. 5B). We also found the pattern of age-specific mortality was very similar in early life for all productivities 𝜅, but that lifespan increased predictably as asymptotic body size increased (Suppl. Fig. 5).

**Figure 5.**
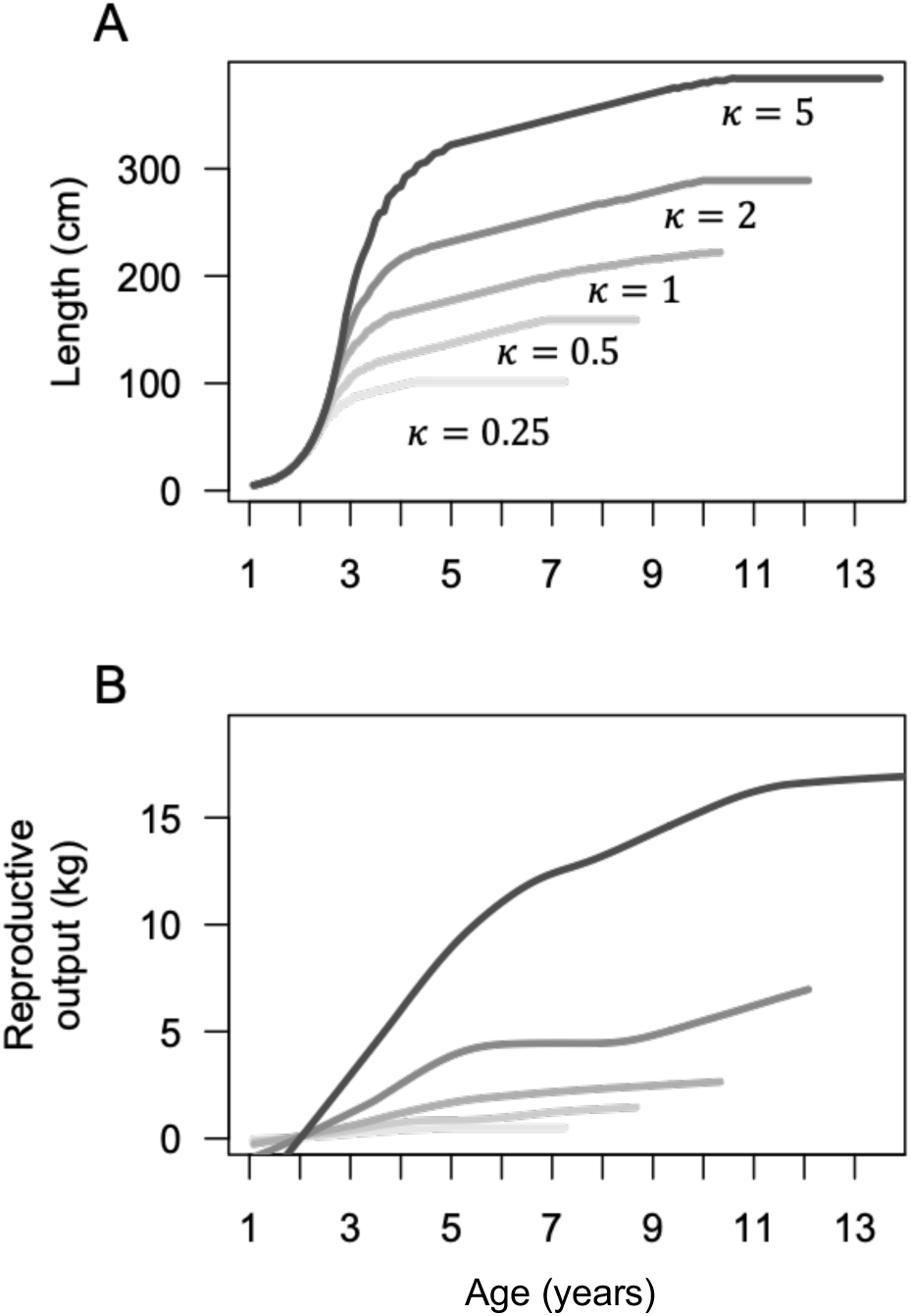
(A) Growth (length at age) and (B) Expected reproductive output of individuals in different productivity 𝜅 scenarios. Each curve ends at the age (maximum lifespan) at which an individual’s cumulative chance of mortality due to predation or starvation is greater than 97%. Temperature in every scenario was constant at 16.85 ℃, and *h* was constant at 8. The reproductive output was smoothed using the loess tool in R; raw data points can be found in Suppl. Fig. 4.

When we compared the interacting effects of ecosystem productivity (𝜅) and metabolic costs of elevated temperature (𝜏) on patterns of growth and reproductive allocation, we found the effects of temperature-dependent costs on body size and reproductive output were relatively small (Fig. 6). The biggest differences emerged in highly productive environments with dramatic increases in costs associated with higher temperatures. Specifically, when average temperature increased by 10 degrees (e.g., from 11.85 to 21.85 ℃ or from 16.85 to 26.85 ℃), maximum body sizes were 3-10% smaller, with the largest differences occurring when 𝜅 = 5 (specific values in Fig. 6 are reported in Suppl. Table 1). At the same time, these increases in temperature led to concomitant reductions in lifetime reproductive output that ranged from 3-10% (Fig 6B). However, it is important to note that comparing thermal increases in 5-degree intervals only generated minor (3-5%) decreases in growth and reproduction, unless both predators and prey were scarce (𝜅 ≤ 0.5, Suppl. Table 1).

**Figure 6.**
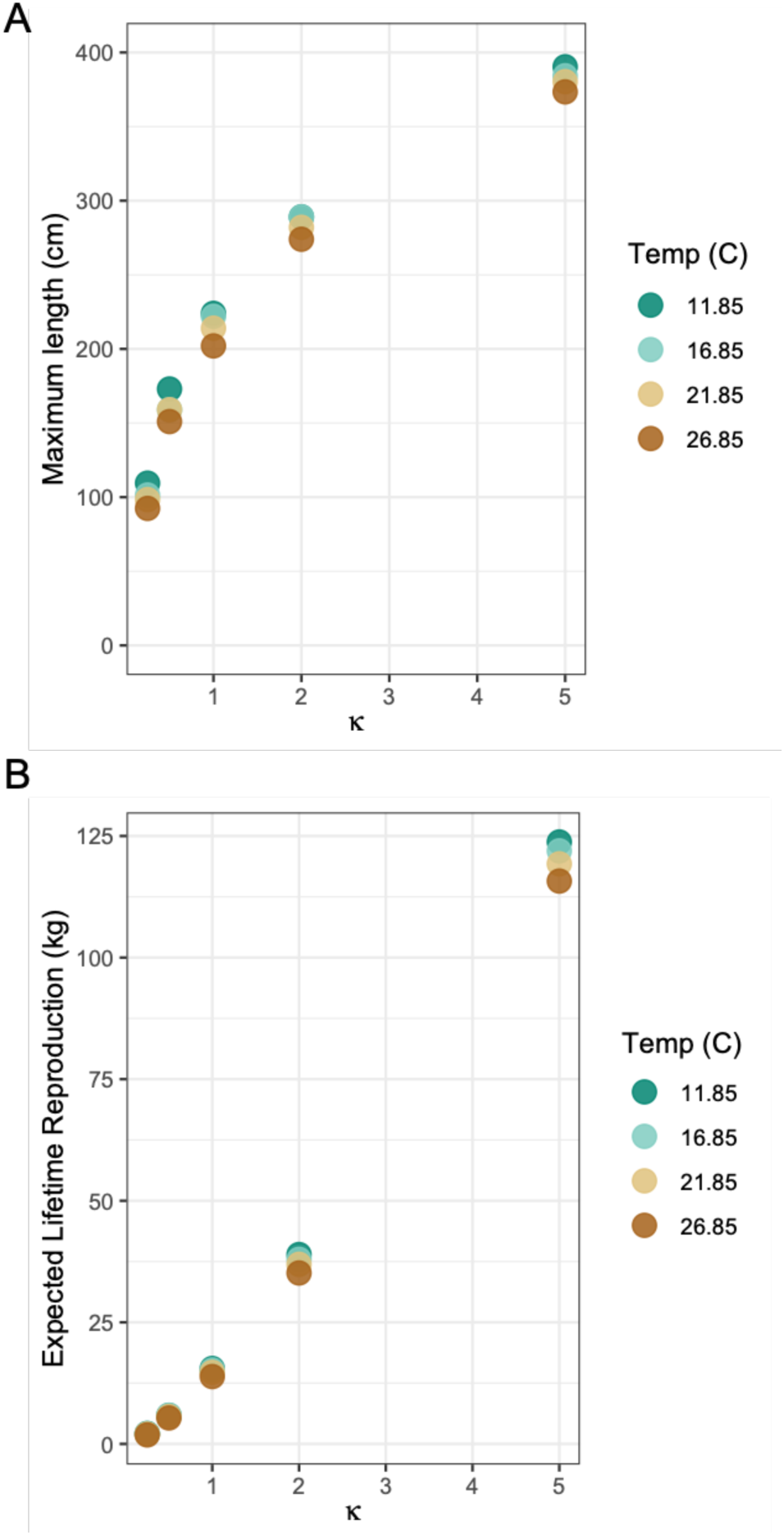
(A) Maximum body size and (B) Expected lifetime reproductive output (in kg) of individuals plotted against a range of productivity scenarios. Color of points corresponds to differences in average annual temperature in℃, representing increased physiological costs.

The interacting effects of temperature and productivity on body size showed that when both prey and predators were abundant, the benefits of growing large outweighed associated increases in metabolic costs. Remaining smaller in thermally costly conditions (to minimize energetic requirements for maintenance) equated to an increased risk of mortality through predation and thus shorter lifespans. In other words, the effect of metabolic costs on lifespan were indirectly expressed through body size.

To understand the robustness of the main results in Figs. 5 and 6, we conducted sensitivity analyses. First, we varied the parameter *h,* representing predator efficiency (Eq. 3; Andersen 2019), to change the allometric properties of the risk of predation (Suppl. Fig. 6). However, while varying this parameter affected maximum lifespan by changing the risk of predation, it did not dramatically change individual growth trajectories or patterns of size-specific reproduction (Suppl. Fig. 6). By contrast, the constraint on the amount of lipid stores that can be spent each month on reproduction, *χπ*, was of more importance for growth because it depended directly on the individual’s structural mass. This parameter contributes biological realism to the state dynamics, because it represents a limit on the maximum possible fat stored as gonadal tissue (i.e., so fish cannot remain small and instead channel all surplus energy toward reproduction with no constraint; at some point their body cavity will limit gonadal tissue). However, it is difficult to measure or estimate directly, especially for fish that spawn multiple times per year, because often little is known about the frequency of spawning. In sensitivity analyses, we found that increased body sizes were favored if we made this constraint more stringent (e.g., decreased *χπ* from 0.2 to 0.1), representing the case that individuals were able to devote less of their body cavity to gonadal tissue. This pattern held at both high and low productivity values (Suppl. Fig. 7).

Finally, we used estimates of parameters for the metabolic cost function that were measured experimentally for tuna species, because of our interest in explaining diversity in tuna growth patterns (Clarke and Johnston 1999). However, to understand if the relatively small effect of increased temperatures on body size in Fig. 6 was robust to differences in metabolic costs, we conducted sensitivity analyses in which the shape and allometric scaling of costs varied (Suppl. Figs. 8 and 9). We confirmed that our choice of parameters for this function had relatively minor influences on growth and reproduction.

In the environmental scenarios representing the three ecological lifestyles of tunas (Table 2) our model predicted growth, body size, and reproductive patterns that were qualitative matches with typical species of each of the three tuna ecological lifestyles (Fig. 7). With the environmental scenario representing the tropical shallow-diving lifestyle, we predicted a maximum body size of 70 cm, longevity of 6-7 years of age, and continuous reproduction at relatively low levels (left column, Fig. 7, Table 2). These predicted traits are similar to those of frigate tuna *Auxis thazard,* a tropical shallow-diving tuna species with small body size. With the environmental scenario parametrized to match the tropical deep-diving lifestyle, we predicted growth to sizes well over 200 cm, lifespans of less than 15 years, and continuous reproductive output that increases over the individual’s life (middle column, Fig. 7, Table 2). While we believe this is broadly consistent with the life histories of tropical deep-diving tuna species such as yellowfin tuna *Thunnus albacares*, lifetime reproductive patterns of these batch-spawning species are not well known (Horswill et al. 2019). In the environmental scenario matching the lifestyle of temperate deep-diving tunas, such as Atlantic bluefin tuna, *Thunnus thunnus,* we predicted the maximum body size exceeded 350 cm a longevity that was longer than 18 years (right column, Fig. 7, Table 2). Spawning was constrained to be seasonal in tunas in the third ecological lifestyle (i.e., spawning could only occur in temperatures above 24 ℃, which we specified individuals experienced for three months of the year).

**Figure 7.**
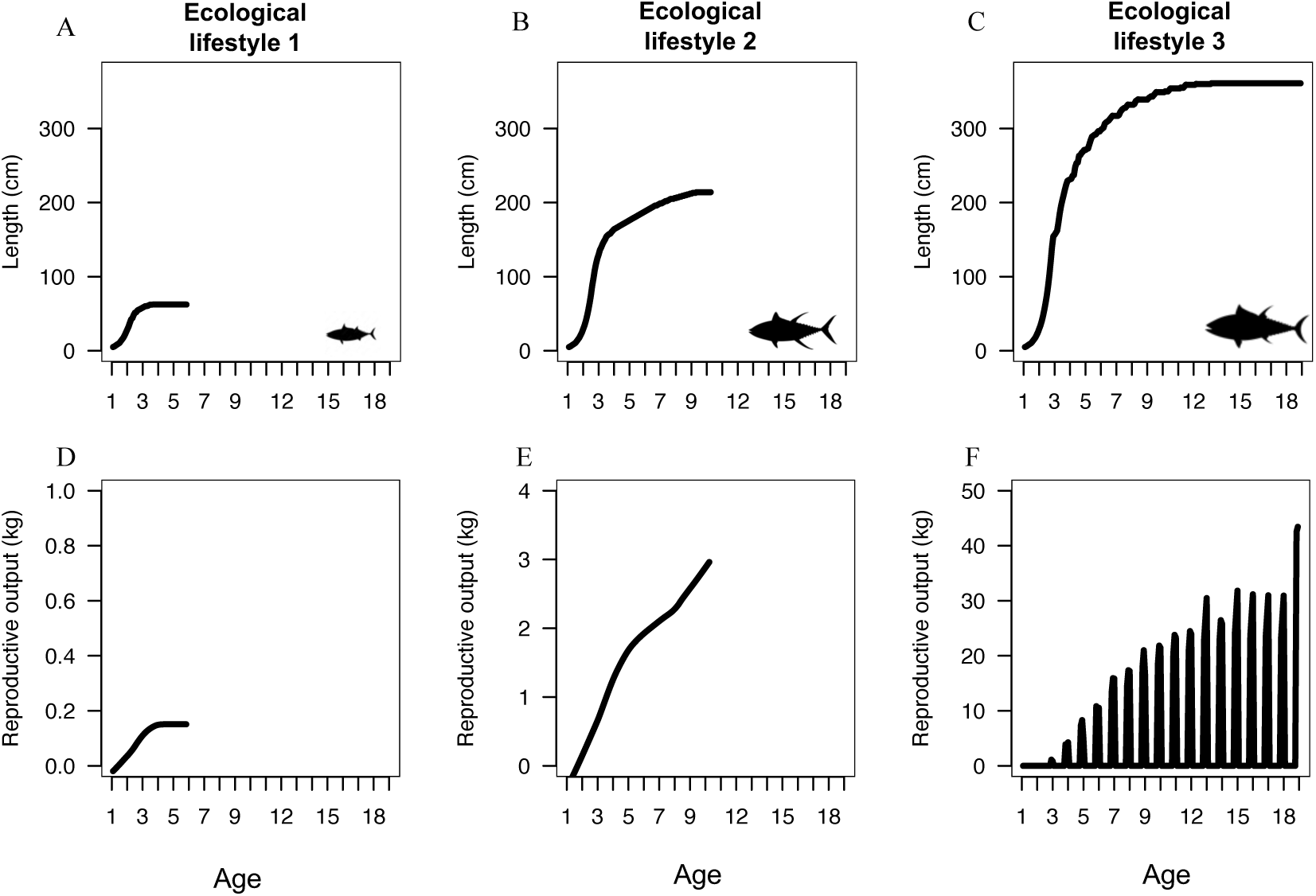
Growth (top row) and reproduction (bottom row) predicted by our general energy-based model for the three characteristic ecological lifestyles of tunas using three representative environmental scenarios (columns). Details of each scenario and species are in Table 2. With the exception of C and F, all curves end at the age when less than 3% of the population is expected to survive, representing the cohort lifespan. A and D represent tropical shallow-diving tuna species such as the frigate tuna *Auxis thazard*. B and E represent tropical deep-diving tuna species such as yellowfin tuna *Thunnus albacares*. C and F represent temperate deep-diving species such as Atlantic bluefin tuna *Thunnus thynnus*, which migrate seasonally from higher latitudes with colder and productive waters to less productive and warmer waters. At age 18, more than 10% of individuals in the temperate deep-diving lifestyle were still alive and are likely to live much longer (C, F). As in Fig. 5, the reproductive output in panels D-E was smoothed using the loess function.

### Analysis of seasonality in temperature and spectrum productivity

The results of the tuna case study yielded an unexpected pattern: when spawning activity, costs associated with increased temperature, and ecosystem productivity varied seasonally (i.e., in the third ecological lifestyle of temperate deep-diving tunas), individuals grew to be substantially larger than the size observed under constant environment conditions that are otherwise comparable in terms of average productivity (𝜅) and temperature (𝜏).

To investigate this pattern further, we made several subsequent comparisons of the effects of seasonality in spawning season length, temperature (in which species spawned for three months in waters that are 9 degrees warmer than their winter foraging grounds) and productivity (which increased threefold during the winter months spent on the foraging grounds). In Fig. 8, for a range of ecosystem productivities (𝜅 = 0.5 - 2.5 when averaged across the 12-month period), we compared the maximum lengths predicted in constant environments with scenarios in which reproductive season length varied (represented by the dash length in Fig. 8), and/or temperature and productivity fluctuated over the year, for a total of 6 contrasting scenarios: (i) the most basic constant scenario when temperature, productivity (𝜅), and spawning were constant year round (solid aqua line in Fig 8, which is identical to the 𝜏 = 16.85 ℃ scenarios shown in Fig. 6); (ii) a seasonal scenario when temperature increased during the three month spawning season but seasonal scenario with a threefold increase in productivity for nine months to represent time spent in foraging grounds, but with temperature kept constant throughout the year (navy blue short-dashed line in Fig. 8); productivity 𝜅 did not change throughout the year (purple short-dashed line in Fig. 8), (iii) a (iv) the full seasonal scenario whesre productivity and temperature varied during the three-month summer when spawning was possible (salmon-pink short-dashed line in Fig 8; note this scenario is similar to Fig. 7C, but with lower values of 𝜅); (v) a full seasonal scenario where the warm, low-productivity season lasted for six months instead of three (yellow long-dashed line in Fig. 8); and (vi) a seasonal scenario where productivity and temperature varied during the three-month summer but spawning was possible during the full twelve months (solid orange line in Fig. 8).

**Figure 8.**
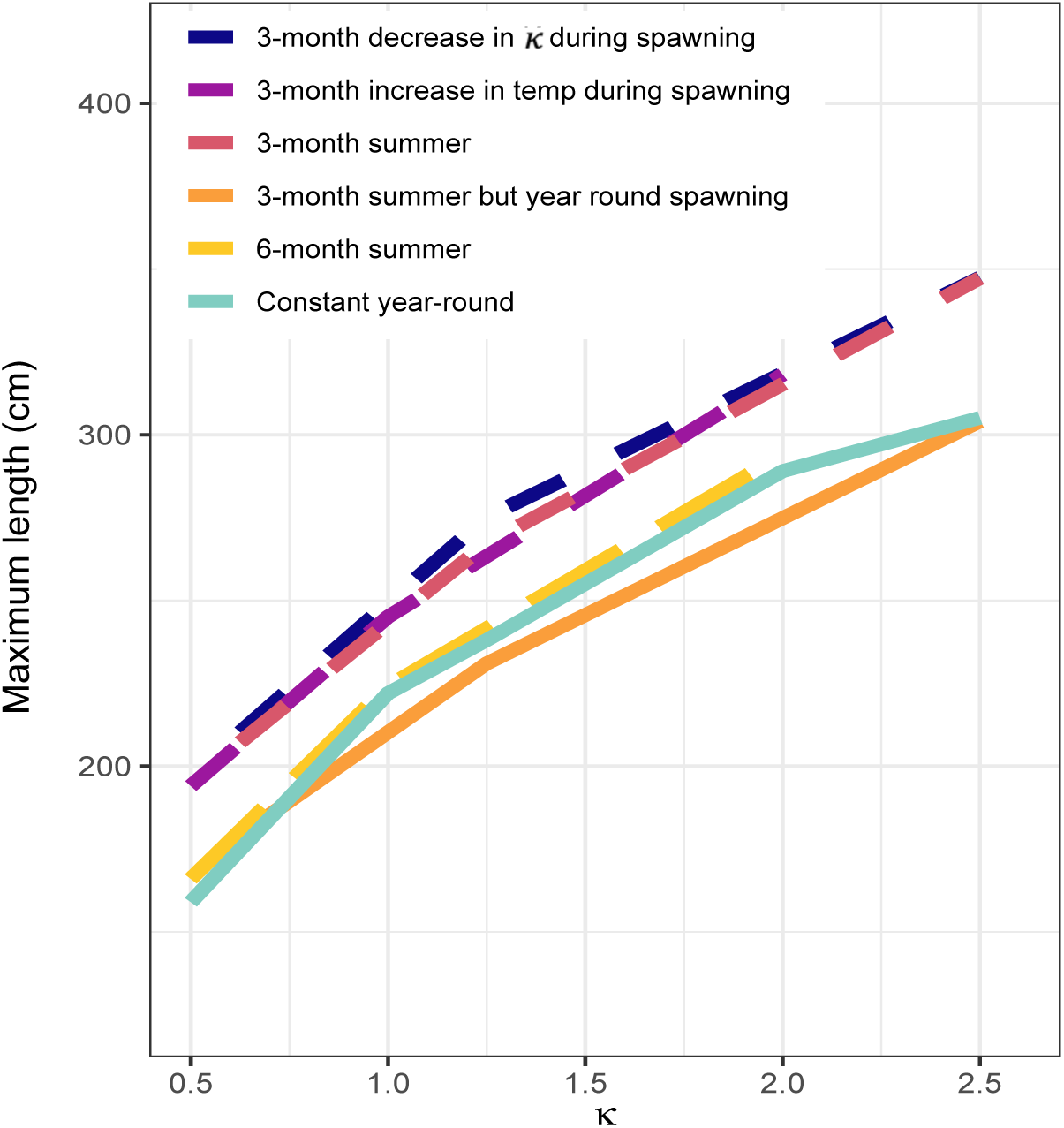
Maximum body length across values of 𝜅 in multiple scenarios varying seasonality in food, temperature, and spawning season duration. As described in the text, solid lines represent scenarios where spawning is possible every month of the year, while the dashed lines represent scenarios where spawning is restricted to three months (short-dashed) and six months (yellow long-dashed) of the year. Colors correspond to different scenarios in which temperature and productivity fluctuate temporally.

These comparisons clarified that the restriction of spawning to three months of the year has the largest effect on maximum body size across a range of ecosystem productivities (because the short-dashed lines are substantially greater than solid or long-dashed lines in Fig. 8). Whenspawning was restricted to three months, the effects of temperature-associated costs and productivity alone were comparable to those when both effects are combined (in Fig. 8, the three short-dashed lines are similar). When spawning was restricted to six months of the year, and temperature and productivity fluctuated seasonally, individuals grew to intermediate sizes.

Smaller sizes were predicted when spawning was possible during every month of the year, especially when temperature and productivity fluctuated seasonally (the solid orange line in Fig. 8). From this set of follow-up analyses, we concluded that resources and temperature effects on energetic budgets were not the primary drivers of larger body sizes in our seasonal model. Rather, the compression of reproductive opportunities into three months of the year, and the temporal separation of reproduction from the most valuable foraging opportunities, favors increased allocation to growth in seasonal environments.

## Discussion

We used state-dependent energetic modeling to test our hypothesis that accounting for trophic interactions represented by size spectra, and the effects of seasonality in metabolic demands and spawning opportunities, can explain observed diversity in life histories of fishes. We then asked whether variation in specific environmental conditions, including seasonality in resources and metabolic costs associated with temperature, can predict the observed life history traits of three tuna species representing different ecological lifestyles. By unpacking the effects of temporal variation in spawning opportunities, metabolic demands, and foraging opportunities, we found that seasonal cycles in resource intake and expenditures have a greater effect on the evolution of body size than that of direct metabolic costs arising from warmer temperatures. Our results do not directly address potential effects of temperature on environmental productivity and individual activity levels, both of which can increase consumption. However, we found that when individuals experienced the same metabolic costs with differing levels of ecosystem productivities, increasing productivity has a positive effect on growth rates. This contrast suggests that net effects of temperature on growth depend on both metabolism and resource availability.

The key difference between our general energetic model and other models of life history evolution of fishes in varying environmental conditions (e.g., Kozłowski and Teriokhin 1999, Lika and Kooijman 2003) is that we consider the mass-specific scaling of metabolic costs with temperature in addition to differences in primary productivity, which drive changes in predator-prey interactions. Across a range of temperatures, our model predicted the benefits of growing large through the size spectrum outweighed the increase in metabolic costs until very large sizes and/or very dramatic increases in temperature. For example, although our model predicted smaller body sizes and decreased lifetime reproduction due to increased costs at higher temperatures when both predators and prey were scarce, these differences were relatively small – all else being equal – until. average temperatures increased by 10-15 ℃. However, increased temperatures may also increase the productivity of ecosystems, offsetting these costs. The parameter representing productivity, 𝜅, had a much stronger influence on our results than any of our other parameters; consistent with expectations, our model predicted larger maximum body sizes, greater reproductive output, and longer lifespans in more productive ecosystems. One exception came when estimating age-specific mortality of each environment, which was more sensitive to parameter values determining predator efficiency in capturing prey (*h,* Eq. 3). While this parameter changed survival, it did not substantially change growth trajectories.

Our results provide a novel explanation for asymptotic growth of fishes. Prior work has predicted asymptotic growth patterns based on physiological arguments for intrinsic geometric constraints (von Bertalanffy 1960, Pauly 2010), or by specifying that increased allocation to reproduction limits growth as organisms age (Jusup et al. 2011). In our model, multiple (mainly extrinsic) ecological mechanisms led to the slowdown of growth and emergence of an asymptotic size in each environment. Unlike previous research, in our results the onset of reproduction alone did not correspond immediately to decreasing growth. Instead, the availability of prey, the physical and temporal limits on resource intake and storage, the risk of predation, and constraints on gonad size and reproductive timing all increased the advantage of growing to larger body sizes; these advantages balanced against metabolic demands and the actual energetic requirements of growth determined maximum body sizes.

We used variation among the ecological lifestyles of tunas to motivate our model parameterizations and comparisons, while also aiming to provide general insights into the evolution of fish life histories. We successfully predicted growth and reproductive patterns that are consistent with species representing each ecological lifestyle of tunas (Fig. 7). We found a previously unknown effect of seasonal variation on body size. Further investigation revealed this effect was robust to fluctuations in metabolic demands or in productivity alone, and instead depended on the spawning season duration. Our findings that large body sizes emerged from the optimal allocation strategy in seasonal environments - whether or not temperature varied - do not support explanations for spatial gradients in body size based on rising metabolic costs with temperature. Instead, our analysis in Fig. 8 supports the conclusion that growth to larger maximum body sizes is driven by the limited time available for reproduction, coupled with the opportunity to forage and store energy in productive ecosystems the rest of the year. These results could explain giantism of fishes in highly seasonal polar environments. Note that the mechanism here contrasts proposed explanations for latitudinal clines in arthropods, which also invoke season length (Blankenhorn and Demont, 2004; Horne et al. 2015), because for these species the growing season is shorter and thus generation time shorter at high latitudes.

Our model illustrates that predicting responses in growth and reproduction in changing environments is complex, because optimal strategies will respond to selective pressures from many factors, including intrinsic constraints on storage and reproduction, physiology, and the nature of ecosystem size spectra. For fishes like tunas, which often exhibit a combination of capital and income breeding, and for which reproduction is constrained by ambient temperature, we expect that as climate change generates warmer conditions, eventually spawning at higher latitudes for longer periods could be possible. However, our results suggest that cascading effects on growth are not easily predicted and will depend on how warming waters reverberate through the size spectrum.

Future work clarifying the complicated relationship between the productivity of oceanic ecosystems and temperature is needed to predict the net effect of changes in climate on the evolution of ectothermic life histories. Changes in climate over recent decades have been shown to affect fish recruitment, growth, and fishery productivity (Free et al. 2019; Oke et al. 2020).

Several mechanisms have been proposed to explain these changes (Fernandes et al. 2020). Early models based on differences in resource intake and metabolic demands predicted that increased temperature will lead to reduced growth and smaller body sizes, i.e., ‘shrinking fishes’ (Cheung et al. 2008), but experimental evidence shows growth is less affected by temperature than reproductive allocation (Wootton et al. 2022), and that metabolic rate can adapt to increased temperatures (Pilakouta et al. 2020). Both physiological models and experimental observations have necessarily minimized confounding effects of environmental productivity, seasonality, and ecological interactions. Our model includes some of the extrinsic abiotic and ecological variables that previous work has had to ignore, but we focused primarily on the effects of temperature on metabolic costs in this framework, and determined that these costs had only minor effects on growth (Fig. 6). Our results imply that the negative effects of temperature on metabolic costs are relatively unimportant compared to the overall productivity of ecosystems, as well as seasonal dynamics of resource acquisition. Yet direct effects of temperature on ecosystem productivity could drive variation in growth and reproduction (Fig. 5). Temperature can change activity levels of both predators and prey, as well as physiological processes such as digestion efficiency.

Including such processes in future investigations of our questions, while increasing model complexity, could be useful to address biological responses to ongoing environmental change. The current model has several additional assumptions that could be explored in future analyses. In the analyses presented here, we have assumed a static relationship between length and mass, and the conversion of mass to joules (energy density of tissue). Keeping these relationships constant ensured a common currency linking individual energetic budgets to ecological changes in foraging success and predation risk. Exploring the consequences of these relationships for differences among clades would be a natural follow-up to our study. Furthermore, while we did not directly address the interaction between of fisheries-induced selection and growth (e.g., Audzijonyte et al. 2016), it would be a rich area of further investigation. Plastic or evolutionary responses to fishing potentially include faster growth, earlier maturation rates, and smaller maximum body sizes, but the consequences for marine food webs are complex (Hočevar and & Kuparinen 2021). The framework introduced here potentially could capture how eco-evolutionary feedbacks between fishing mortality, length and mass relationships, and ecosystem size spectra interactively affect trends in body sizes of focal species. Studies of the interaction between fishing and size spectra have been focused on lake systems where food webs have been studied in greater detail (Perälä and Kuparinen 2020). Understanding how these physiological and ecological processes are mechanistically linked is necessary to understand how species will respond to different environmental conditions in future oceans.

In summary, our findings suggest that when predicting future growth patterns under projected changes in climate, ecological and physiological factors - primary production, prey size and availability, and predation risk - will all play a greater role than metabolic demands in determining trends in maximum body size. Nevertheless, we offer our model as a step toward models of fish growth with higher fidelity to nature that incorporate not only physiology and energetic allocation budgets but are also embedded in ecosystem size spectra. We hope that further exploration of our approach may lead to reconciliation of divergent results regarding the effect of climate change on fish body sizes.

## Appendix. Optimization details

Dynamic programming equations (Eq. 12) are constrained optimization algorithms with the purpose of determining the optimal set of behaviors or decisions that maximize a quantitative metric over time, such as lifetime expected reproductive output (fitness). A key property is that the decisions at one time affect the state variables at the next time. When solving Eq. 12, we consider how the allocation decisions an individual makes during one month of its life affect its future size, energetic reserves (lipid stores) and chances of mortality, and find the set of allocation to growth and reproduction that maximizes its fitness in a given environment. We are able to calculate lifetime fitness by solving the equation using backwards iteration. By starting at 𝑡 = 𝑇 – 1, and assuming there is no possibility of future fitness (i.e., the second term of Eq. 12 is equal to zero for all possible values of length and lipid stores) we can populate the array with the fitness of all possible combinations of length, lipid stores, and allocation strategies at 𝑡 = 𝑇 – 1; this fitness will be the first term on the right side of Eq. 12. This process is repeated, working backwards, until *t =* 1. We can thereby determine the combination of *g* and *r* that maximizes both current fitness at *t*, and the fitness expected from 𝑡 + 1 until *T*, given the emergent chances of survival (which varies due to both the risk of predation and starvation), length, and lipid stores resulting from that particular strategy. Solving Eq. 12 in this way produces an array storing the proportional allocation to growth and reproduction that leads to the highest lifetime fitness, for all possible combinations of size and lipid stores (𝑙 and 𝑠) for every month until the final time *T*.

Solving a dynamic programming equation with two state variables, such as Eq. 12, is computationally expensive, and we employed a number of techniques to make the iteration more efficient and to approximate a smooth fitness surface. First, we constrained the parameter space that was evaluated. Specifically, we used an integer index *I* (with a maximum of 𝐼*_max_*) to represent lipid stores *s,* converting the index to values in joules in the dynamic loop. We then related the range of *s* we explored to each value of *l* (because lipid stores are constrained by *l*), by setting

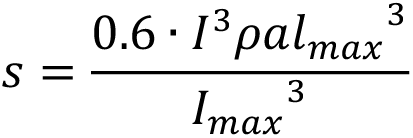

In other words, we adjusted the numerical step size for possible values of *s* to be finer for smaller individuals. Furthermore, we used linear interpolation of state values (Clark and Mangel 2000) when computing expected future fitness in Eq. 12 to minimize discontinuities on the fitness landscape arising from the integer index of energetic state. We did not interpolate length, since its unit (centimeters) was sufficiently fine-grained that there were minimal effects of discontinuities.

We found that sharp transitions in the fitness landscape, such as at threshold values representing constraints, hindered our optimization algorithm. Therefore, we used an asymptotic function for the fitness increment associated with a given level of reproductive output. We let 𝑅(𝑟, 𝑠, 𝑙, 𝑡) denote the potential increment in fitness in month *t* when 𝐿(𝑡) = 𝑙, 𝑆(𝑡) = 𝑠, and a fraction of stores *r* is allocated to current reproduction, and assumed that 𝑅 increased smoothly toward the maximum possible for the given length *l:*

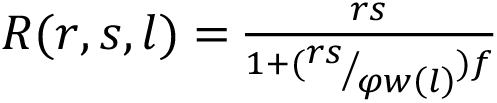

The asymptotic value of this function depends on the value of *f* in the denominator, which controls the abruptness of the constraint on current fitness (Suppl. Fig. 2). In other words, the steepness of the multivariate landscape around the fitness optimum is modulated by *f.* For simplicity, we assume 𝑓 = 1 for all results presented hereafter. In that case, when 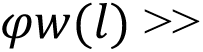 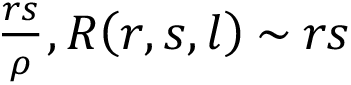 and when 𝑟𝑠 >> 𝜑𝑤(𝑙), then 𝑅(𝑟, 𝑠, 𝑙) ∼ 𝜑𝑤(𝑙).

All programming was done using R 4.1.1. We ran the code to solve the dynamic programming equation and simulate the individual life histories using Rscript commands from the Linux shell of a 2019 MacPro with 16 cores and 96 GB of RAM. Jobs were run in parallel and the runtime of each job was between 60 and 120 minutes. The results of each job are presented in the figures summarizing growth, reproduction, and survival data; these analyses and figures were produced in the Rstudio IDE.

## Acknowledgements

This work was supported by NSF-NERC DEB-1556779 to HKK and JM, and NSF DEB 15-55729 and ONR N000141912494 to MM. CH is funded by Research England. NKD was supported by the Discovery and Accelerator grants from Natural Science and Engineering Research Council and the Canada Research Chair program. We thank K. H. Andersen for discussion of early drafts and J. A. Draghi for help troubleshooting the optimization.

## Data Availability Statement

The data and code that support the findings of this study are openly available in an archived Github repository available at https://zenodo.org/records/8102868.

## Data Archiving statement

All code and model outputs for this study are available in a github repository and will be archived upon publication.

## References

Álvarez-Noriega M., White C.R., Kozłowski J., Day T., Marshall D.J. 2023. Life history optimisation drives latitudinal gradients and responses to global change in marine fishes. PLoS Biol 21(5): e3002114.

Andersen, K. H. 2019. Fish Ecology, Evolution, and Exploitation: A New Theoretical Synthesis. Princeton University Press, Princeton.

Andersen, K. H., Jacobsen, N. S., Farnsworth, K. D., and J. Baum. 2016. The theoretical foundations for size spectrum models of fish communities. Canadian Journal of Fisheries and Aquatic Sciences 73:575–588.

Audzijonyte, A., Fulton, E., Haddon, M., Helidoniotis, F., Hobday, A.J., Kuparinen, A., Morrongiello, J., Smith, A.D., Upston, J. and Waples, R.S., 2016. Trends and management implications of human-influenced life-history changes in marine ectotherms. Fish and Fisheries, 17(4), pp.1005–1028.

Audzijonyte, A., Richards, S.A., Stuart-Smith, R.D., Pecl, G., Edgar, G.J., Barrett, N.S., Payne, N. and Blanchard, J.L., 2020. Fish body sizes change with temperature but not all species shrink with warming. Nature Ecology & Evolution. 4:809–814.

Audzijonyte, A. and Richards, S.A., 2018. The energetic cost of reproduction and its effect on optimal life-history strategies. American Naturalist. 192:E150–E162.

Audzijonyte, A., Jakubavičiūtė, E., Lindmark, M., and Richards S.A. 2022. Mechanistic Temperature-Size Rule Explanation Should Reconcile Physiological and Mortality Responses to Temperature. The Biological Bulletin 2022 243:2, 220-238

Barnes, W., Maxwell, D., Reuman, D.C., and Jennings, S., 2010. Global patterns in predator-prey size relationships reveal size dependency of trophic transfer efficiency. Ecology 91:222–232.

Benoît, E., and M. J. Rochet. 2004. A continuous model of biomass size spectra governed by predation and the effects of fishing on them. Journal of Theoretical Biology 226:9–21.

Bernal, D. 2011. Physiological specializations of different fish groups. Pelagic fishes. Pages 1887-1902 in Farrell AP (ed) Encyclopedia of Fish Physiology: From Genome to Environment, Vol 3. Academic Press, San Diego.

Bernal, D., Sepulveda C., Musyl, M., Brill, R. (2009) The eco-physiology of swimming and movement patterns of tunas, billfishes, and large pelagic sharks. Pages 436–483 in Domenici P, Kapoor BG (eds) Fish Locomotion: an Etho-Ecological Approach. Science Publishers, Enfield.

Bernal, D., Brill, R.W., Dickson, K.A. and H.A. Shiels. 2017. Sharing the water column: physiological mechanisms underlying species-specific habitat use in tunas. Reviews in Fish Biology and Fisheries 27:843–880.

Bernhardt, J.R., Sunday, J.M. and M.I. O’Connor. 2018. Metabolic theory and the temperature-size rule explain the temperature dependence of population carrying capacity. American Naturalist. 192:687–697.

Beverton, R. J. H., and S. J. Holt. 1959. A review of the lifespans and mortality rates of fish in nature, and their relation to growth and other physiological characteristics. Pages 142–177 *in* CIBA Foundation Colloquium on Ageing.

Bigman, J.S., M’Gonigle, L., Wegner, N.C. and Dulvy, N.K. 2021. Respiratory capacity is twice as important as temperature in driving patterns of metabolic rate across the vertebrate tree of life. Science Advances 7: eabe5163.

Blanchard, J. L., Heneghan, R. F., Everett, J. D., Trebilco, R. and A. J. Richardson. 2017. From bacteria to whales: using functional size spectra to model marine ecosystems. Trends in Ecology & Evolution. 32:174–186.

Blanchard, J. L., Jennings, S., Law, R., Castle, M.D., McCloghrie, P., Rochet, M.J., and E. Benoît. 2009. How does abundance scale with body size in coupled size-structured food webs? Journal of Animal Ecology. 78:270–280.

Blanchard, J.L., Jennings, S., Holmes, R., Harle, J., Merino, G., Allen, J.I., Holt, J., Dulvy, N.K. and M. Barange. 2012. Potential consequences of climate change for primary production and fish production in large marine ecosystems. Phil. Trans. Roy. Soc. B 367:2979–2989.

Blanckenhorn, W.U. & Demont, M. 2004. Bergmann and converse Bergmann latitudinal clines in Arthropods: two ends of a continuum? Integr. Comp. Biol., 44, 413–424

Brown, J. H., J. F. Gillooly, A. P. Allen, V. M. Savage, and G. B. West. 2004. Toward a metabolic theory of ecology. Ecology. 85:1771–1789.

Chapman, E. W., Kørgensen, C., and M. E. Lutcavage. 2011. Atlantic bluefin tuna (*Thunnus thynnus*): a state-dependent energy allocation model for growth, maturation, and reproductive investment. Canadian Journal of Fisheries and Aquatic Sciences 68:1934–1951.

Charnov, E. L., Berrigan, D., and R. J. H. Beverton. 1991. Dimensionless numbers and the assembly rules for life histories. Phil. Trans. Roy. Soc. B 332:41–48.

Charnov, E. L., and J. R. Krebs. 1974. On clutch-size and fitness. Ibis. 116:217–219.

Cheung, W.W., Close, C., Lam, V., Watson, R., and D. Pauly 2008. Application of macroecological theory to predict effects of climate change on global fisheries potential. Marine Ecology Progress Series. 365:187–197.

Christensen, A. and Andersen, K.H., 2011. General classification of maturation reaction-norm shape from size-based processes. Bulletin of Mathematical Biology, 73(5): 1004–1027.

Cichoń, M. and Kozłowski, J., 2000. Ageing and typical survivorship curves result from optimal resource allocation. Evolutionary Ecology Research, 2(7): 857–870.

Clark, C. W., and M. Mangel. 2000. Dynamic state variable models in ecology: methods and applications. Oxford University Press, Oxford.

Clarke, A. 2006. Temperature and the metabolic theory of ecology. Functional Ecology 20:405–412.

Clarke, A., and N. M. Johnston. 1999. Scaling of metabolic rate with body mass and temperature in teleost fish. Journal of Animal Ecology. 68: 893–905.

Collette, B.B., Reeb, C. and Block, B.A., 2001. Systematics of the tunas and mackerels (Scombridae). Fish Physiology, 19, pp.1–33.

Conover, D. O., and S. B. Munch. 2002. Sustaining fisheries yields over evolutionary timescales. Science 297:94–96.

Daufresne, M, Lengfellner, K., and U. Sommer. 2009. Global warming benefits the small in aquatic ecosystems. Proc. Nat. Acad. Sci. 106:12788–12793.

Dickson, K.A., and Graham, J.B. 2004. Evolution and consequences of endothermy in fishes Physiological and Biochemical Zoology 77: 998–1018.

Ejsmond, M.J., Czarnołęski, M., Kapustka, F. and Kozłowski, J., 2010. How to time growth and reproduction during the vegetative season: an evolutionary choice for indeterminate growers in seasonal environments. The American Naturalist 175(5): 551–563.

Fernandes, J.A., Rutterford, L., Simpson, S.D., Butenschön, M., Frölicher, T.L., Yool, A., Cheung, W.W. and Grant, A., 2020. Can we project changes in fish abundance and distribution in response to climate? Global Change Biology. 26:3891–3905.

Free, C.M., Thorson, J.T., Pinsky, M.L., Oken, K.L., Wiedenmann, J. and Jensen, O.P. 2019. Impacts of historical warming on marine fisheries production. Science. 363: 979–983.

Froese R. 2006. Cube law, condition factor and weight-length relationships: history, meta-analysis and recommendations. Journal of Applied Ichthyology 22: 241–253

Gadgil, M., and W. H. Bossert. 1970. Life historical consequences of natural selection. American Naturalist 104:1–24.

Giacomini, H.C., Shuter, B.J. and Lester, N.P., 2013. Predator bioenergetics and the prey size spectrum: Do foraging costs determine fish production? Journal of Theoretical Biology 332: 249–260.

Gillooly, J. F., Brown, J. H., West, G. B., Savage, V. M., and E. L. Charnov. 2001. Effects of size and temperature on metabolic rate. Science. 293:2248–2251.

Gislason, H., Daan, N. Rice, J. C., and J. G. Pope. 2010. Size, growth, temperature and the natural mortality of marine fish. Fish and Fisheries 11:149–158.

Hilborn, R. and Mangel, M. 1997. The Ecological Detective. Princeton University Press, Princeton, NJ.

Hočevar, S., & Kuparinen, A. (2021). Marine food web perspective to fisheries-induced evolution. Evolutionary Applications, 14, 2378– 2391.

Horne, C.R., Hirst, A.G., & D. Atkinson. 2015. Temperature-size responses match latitudinal-size clines in arthropods, revealing critical differences between aquatic and terrestrial species. Ecology Letters 18: 327–335

Horswill, C., H. K. Kindsvater, M. J. Juan-Jordá, N. K. Dulvy, M. Mangel, and J. Matthiopoulos. 2019. Global reconstruction of life-history strategies: A case study using tunas. Journal of Applied Ecology 56:855–865.

Houston, A., C. Clark, J. McNamara, and M. Mangel. 1988. Dynamic Models in Behavioural and Evolutionary Ecology. Nature. 332:29–34

Houston, A., and J. McNamara. 1999. Models of Adaptive Behaviour. Cambridge University Press.

Hulthén, K., Hill, J.S., Jenkins, M.R. and Langerhans, R.B., 2021. Predation and resource availability interact to drive life-history evolution in an adaptive radiation of livebearing fish. Frontiers in Ecology and Evolution, 9, p.619277.

Jørgensen, C., K. Enberg, and M. Mangel. 2016. Modelling and interpreting fish bioenergetics: a role for behaviour, life-history traits and survival trade-offs. Journal of Fish Biology 88:389– 402.

Jørgensen, C., and Ø. Fiksen. 2006. State-dependent energy allocation in cod (*Gadus morhua*). Canadian Journal of Fisheries and Aquatic Sciences 63:186–199.

Juan-Jordá, M. J., I. Mosqueira, J. Freire, and N. K. Dulvy. 2013. The Conservation and Management of Tunas and Their Relatives: Setting Life History Research Priorities. PLoS ONE 8:e70405.

Juan-Jordá, M.J., Mosqueira, I., Freire, J., Ferrer-Jordá, E. and Dulvy, N.K. 2016. Global scombrid life history data set. Ecology, 97: 809–809.

Jusup, M., Klanjscek, T., Matsuda, H., Kooijman, S.A.L.M. 2011. A Full Lifecycle Bioenergetic Model for Bluefin Tuna. PLoS ONE 6(7): e21903.

Kindsvater, H.K., Alonzo, S.H., Mangel, M. and Bonsall, M.B., 2010. Effects of age-and state-dependent allocation on offspring size and number. Evolutionary Ecology Research, 12(3), pp.327–346.

Kingsolver, J. G., and R. B. Huey. 2008. Size, temperature, and fitness: three rules. Evolutionary Ecology Research 10:251–268.

Kiørboe, T., and A. G. Hirst. 2014. Shifts in mass scaling of respiration, feeding, and growth rates across life-form transitions in marine pelagic organisms. American Naturalist 183:E118–E130.

Kitchell, J.F., Neill, W.H., Dizon, A.E., and J.J. Magnuson. 1978. Bioenergetic Spectra of Skipjack and Yellofin Tunas. In:Sharp, G.D., Dizon, A.A. (Eds.), The Physiological Ecology of Tunas. Academic Press.

Kozłowski, J., 1992. Optimal allocation of resources to growth and reproduction: implications for age and size at maturity. Trends in Ecology and Evolution, 7(1):15–19.

Kozlowski, J. 1996. Optimal allocation of resources explains interspecific life-history patterns in animals with indeterminate growth *Proc*. R. Soc. B. 263: 559–566.

Kozłowski, J. and Teriokhin, A.T., 1999. Allocation of energy between growth and reproduction: the Pontryagin Maximum Principle solution for the case of age-and season-dependent mortality. Evolutionary Ecology Research 1(4): 423–441

Law, R., M. J. Plank, A. James, and J. L. Blanchard. 2009. Size-spectra dynamics from stochastic predation and growth of individuals. Ecology 90:802–811.

Lika, K. and Kooijman, S.A., 2003. Life history implications of allocation to growth versus reproduction in dynamic energy budgets. Bulletin of Mathematical Biology, 65(5), pp.809–834.

Lindmark, M., Ohlberger, J., & Gårdmark, A. (2022). Optimum growth temperature declines with body size within fish species. Global Change Biology, 28, 2259–271.

Oke, K.B., Cunningham, C.J., Westley, P.A.H., Baskett, M.L., Carlson, S.M., Clark, J., Hendry, A.P., Karatayev, V.A., Kendall, N.W., Kibele, J., Kindsvater, H.K., Kobayashi, K.M., Lewis, B., Munch, S., Reynolds, J.D., Vick, G.D., and E.P. Palkovacs. 2020. Recent declines in salmon body size impact ecosystems and fisheries. Nature communications, 11:1–13.

Perälä, T, Kuparinen, A. Eco-evolutionary dynamics driven by fishing: From single species models to dynamic evolution within complex food webs. Evol Appl. 2020; 13: 2507– 2520.

Perrin, N. and Sibly, R.M., 1993. Dynamic models of energy allocation and investment. Annual Review of Ecology and Systematics, 24: 379–410.

Perrin, N., Sibly, R.M. and Nichols, N.K., 1993. Optimal growth strategies when mortality and production rates are size-dependent. Evolutionary Ecology, 7(6), pp.576–592.

Mangel, M. 2006. The Theoretical Biologist’s Toolbox : Quantitative Methods for Ecology and Evolutionary Biology. Cambridge University Press.

Mangel, M. 2015. Stochastic dynamic programming illuminates the link between environment, physiology, and evolution. Bulletin of Mathematical Biology 77:857–877.

Mangel, M. and C. W. Clark. 1988. Dynamic Modelling in Behavioral Ecology. Princeton University Press, Princeton.

Monsch, K.A. 2000 The Phylogeny of the Scombroid Fishes. PhD thesis, University of Bristol p 272.

Neubauer, P., and K. H. Andersen. 2019. Thermal performance of fish is explained by an interplay between physiology, behaviour and ecology. Conservation Physiology 7:coz025

Pauly D. 2010. Gasping fish and panting squids: oxygen, temperature and the growth of water-breathing animals. International Ecology Institute, Germany.

Pilakouta, N, Killen, SS, Kristjánsson, BK, et al. 2020. Multigenerational exposure to elevated temperatures leads to a reduction in standard metabolic rate in the wild. Funct Ecol. 34:1205–1214.

Pignalosa, P., Pappalardo, L., Gioacchini, G., Carnevali, O. 2020. Length-weight relationships and a new length conversion factor for Atlantic Bluefin tuna (*Thunnus thynnus L*.) caught in the Mediterranean Sea. Collect. Vol. Sci. Pap. ICCAT, 76(2): 138-146.

Quince, C., Abrams, P.A., Shuter, B.J. and Lester, N.P., 2008. Biphasic growth in fish I: theoretical foundations. Journal of Theoretical Biology, 254(2): 197–206.

Schaefer, K.M. 2001. Reproductive biology of tunas. Fish Physiology. 19: 225–270.

Schaefer K.M., Fuller D.W., and B.A. Block 2009. Vertical movements and habitat utilization of skipjack (Katsuwonus pelamis), yellowfin (Thunnus albacares), and bigeye (Thunnus obesus) tunas in the Equatorial Eastern Pacific Ocean, ascertained through archival tag data. Pages 121-144 in Nielsen, J.L., Arrizabalaga, H., Fragoso, N., Hobday, A., Lutcavage, M., Sibert, J. (Eds.), Tagging and Tracking of Marine Animals with Electronic Devices. Springer, New York.

Sheldon, R.W., Prakash, A. and Sutcliffe Jr., W.H. 1972. The distribution of particles in the ocean. Limnology and Oceanography. 17:327–340.

Sheldon, R.W., Sutcliffe Jr., W.H., and M.A. Paranjape. 1977. Structure of pelagic food chain and relationship between plankton and fish production. Journal of the Fisheries Research Board of Canada. 34: 2344–2353.

Shuter, B.J., Giacomini, H.C., de Kerckhove, D. and Vascotto, K., 2016. Fish life history dynamics: shifts in prey size structure evoke shifts in predator maturation traits. Canadian Journal of Fisheries and Aquatic Sciences. 73(4): 693–708.

Southwood, T.R.E. 1977. Habitat, the Templet for Ecological Strategies? Journal of Animal Ecology. 46:337–365.

Sprules, W. G., and L. E. Barth. 2016. Surfing the biomass size spectrum: some remarks on history, theory, and application. Canadian Journal of Fisheries and Aquatic Sciences 73:477–495.

Thunell, V., Gårdmark, A., Huss, M., and Vindenes, Y. 2023. Optimal energy allocation trade-off driven by size-dependent physiological and demographic responses to warming.” Ecology 104(4): e3967.

Thygesen, U.H., Farnsworth, K.D., Andersen, K.H., Beyer, J.E. 2005. How optimal life history changes with the community size-spectrum. Proc. R. Soc. B. 272:1323–1331.

Trebilco, R., Baum, J. K., Salomon, A. K., and N. K. Dulvy. 2013. Ecosystem ecology: Size-based constraints on the pyramids of life. Trends in Ecology & Evolution. 28:423–431.

Ursin, E. 1973. On the prey size preferences of cod and dab. Meddelelser fra Danmarks Fiskeriog Havundersøgelser 7: 85–98.

Varpe, Ø., 2017. Life history adaptations to seasonality. Integrative and comparative biology, 57(5), pp.943–960.

von Bertalanffy L. 1960. Principles and theory of growth. Pages 137–259 *in* Nowinski, W.W., Ed., Fundamental Aspects of Normal and Malignant Growth, Elsevier, Amsterdam.

Walsh, M. R., and D. N. Reznick. 2009. Phenotypic diversification across an environmental gradient: a role for predators and resource availability on the evolution of life histories. Evolution 63:3201–3213.

White, C.R., Alton, L.A., Bywater, C.L., Lombardi, E.J. and Marshall, D.J., 2022. Metabolic scaling is the product of life-history optimization. Science, 377(6608), pp.834–839.

Wong, S., Bigman, J.S., and N.K. Dulvy. 2021. The metabolic pace of life histories across fishes. Proc. R. Soc. B. 288:20210910.

Wootton, H.F., Morrongiello, J.R., Schmitt, T. & Audzijonyte, A. 2022. Smaller adult fish size in warmer water is not explained by elevated metabolism. Ecology Letters, 25, 1177– 1188.

Yanco, S.W., Pierce, A.K. and Wunder, M.B. 2022. Life history diversity in terrestrial animals is associated with metabolic response to seasonally fluctuating resources. Ecography. 2022:e05900

