## Supplementary Tables and Figures for "Size-dependence of food intake and mortality interact with temperature and seasonality to drive diversity in fish life histories"

Supplementary Material for **Fear and foraging interact with temperature and seasonality to drive diversity in fish life histories**

**Supplemental Table 1.** Model outputs corresponding to points in main text Figure 6. All scenarios had constant environments.

| **Temperature $\tau$ in ℃** | $\kappa$ | **Lifetime reproductive output (kg)** | **Maximum body length (cm)** |
|---|---|---|---|
| 11.85 | 0.25 | 2.15 | 109.5 |
| 11.85 | 0.5 | 5.87 | 173 |
| 11.85 | 1 | 15.51 | 224 |
| 11.85 | 2 | 38.94 | 289.1 |
| 11.85 | 5 | 123.77 | 390.4 |
| 16.85 | 0.25 | 2.08 | 101.5 |
| 16.85 | 0.5 | 5.9 | 159 |
| 16.85 | 1 | 15.2 | 222 |
| 16.85 | 2 | 37.97 | 289 |
| 16.85 | 5 | 121.95 | 384.3 |
| 21.85 | 0.25 | 2.02 | 98.4 |
| 21.85 | 0.5 | 5.65 | 159 |
| 21.85 | 1 | 14.73 | 214 |
| 21.85 | 2 | 36.93 | 282 |
| 21.85 | 5 | 119.28 | 380.3 |
| 26.85 | 0.25 | 1.85 | 92.4 |
| 26.85 | 0.5 | 5.33 | 151 |
| 26.85 | 1 | 13.85 | 202 |
| 26.85 | 2 | 35.15 | 274 |
| 26.85 | 5 | 115.73 | 373.4 |

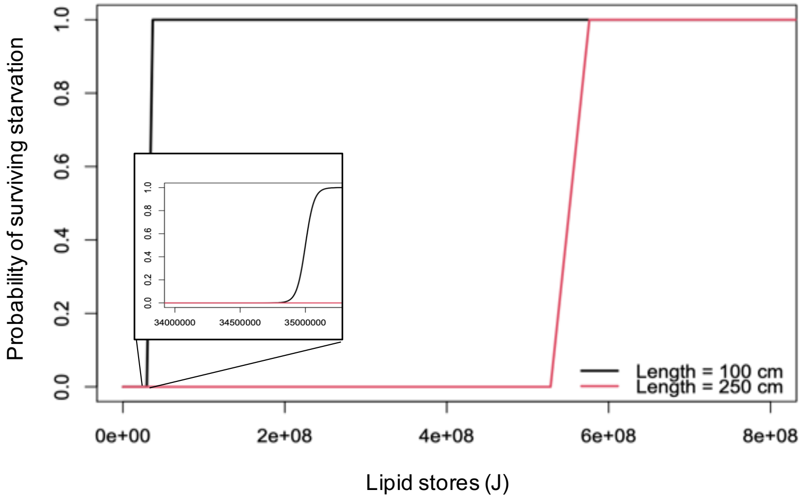


**Supplementary Figure 1**. Main panel: The sigmoid relationship between lipid stores (in joules) and survival for individuals of different lengths. The threshold amount of energy required for metabolic maintenance is a fixed proportion of body mass. The probability of avoiding starvation $\gamma_s(s, l)$ is a logistic function of mass given in Eq. 11. Inset: closeup of the x-axis in the region encompassing the smaller threshold.

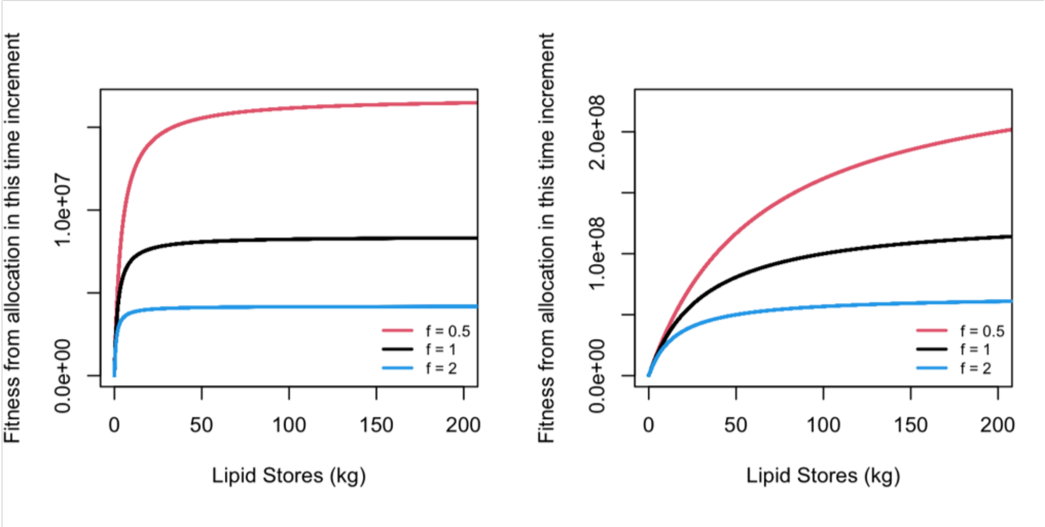


**Supplementary Figure 2**. The fitness increment expected with different amounts of lipid reserves (energy stores) Left panel: Fitness increment for an individual of 100 cm. Right panel: Fitness increment for an individual of 250 cm. We show the shape of the function for three values of *f* but for all analyses here we used *f* = 1.

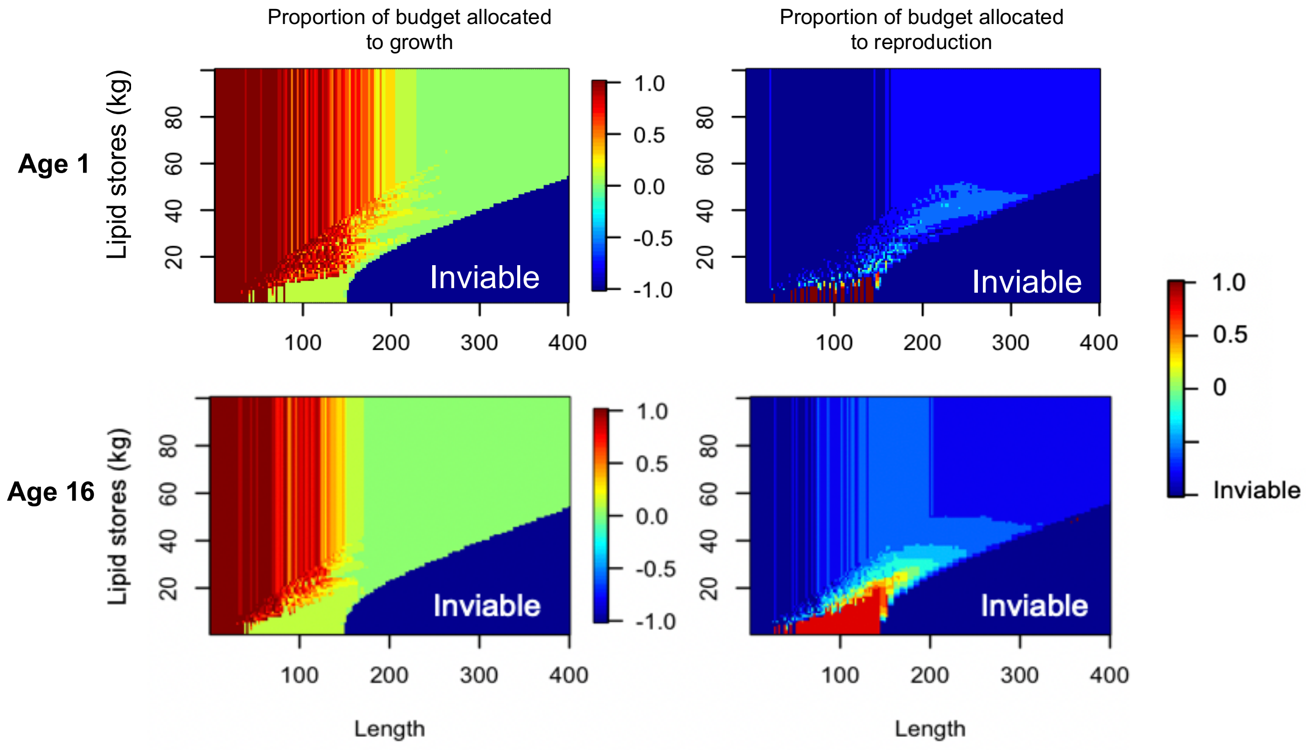


**Supplementary Figure 3.** Example rules for allocation to growth and reproduction for ages 1 and 16. Colors represent the proportional allocation of stores to growth (left column) or reproduction (right column) that is optimal for a given combination of state and length. The inviable region (dark blue) represents parameter space where remaining stores are insufficient for survival, given the individual's length (in other words, energetic requirements for maintenance are not met). Comparing panels within rows shows differences between allocation to growth and reproduction as a function of state and length. Comparing panels within columns shows how the tradeoff between growth and reproduction changes with age (here, reported in years). Note that the striations come from the discretization of allocation values of $g$ and $r$ in Eq. 12.

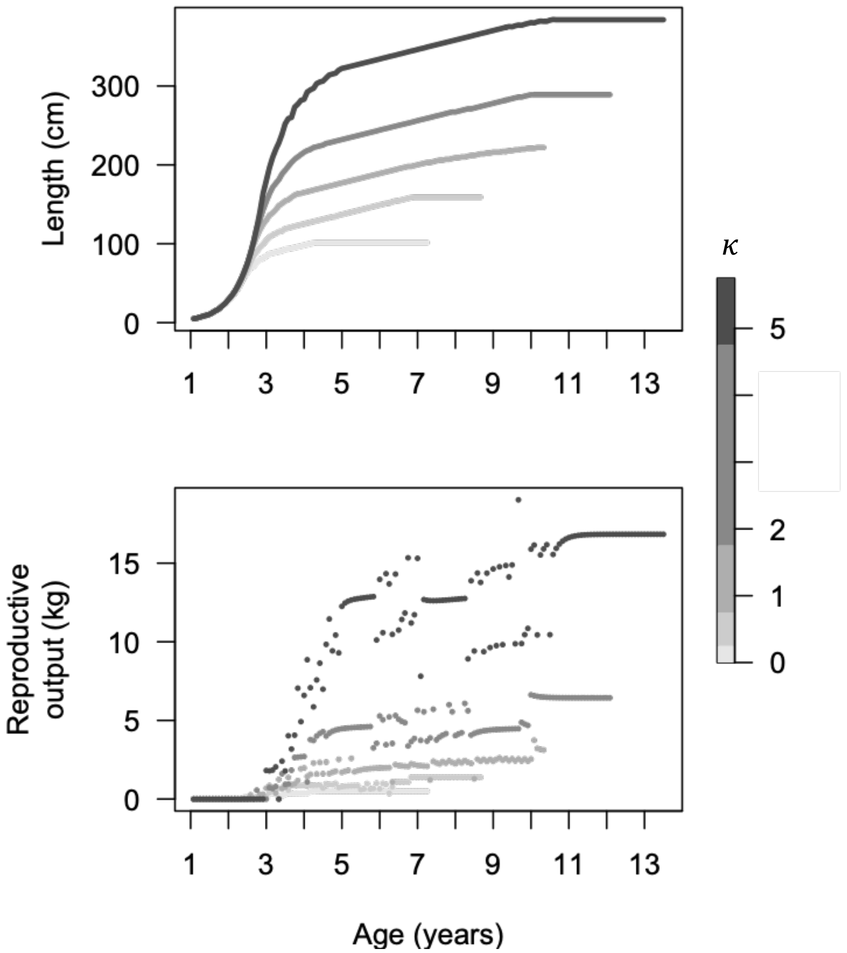


**Supplementary Figure 4.** Top: The emergent pattern of growth at different levels of $\kappa$ (this panel is identical to Fig. 4A in the main text). Bottom: The raw points from Fig. 4B in the main text (which presents the loess curve).

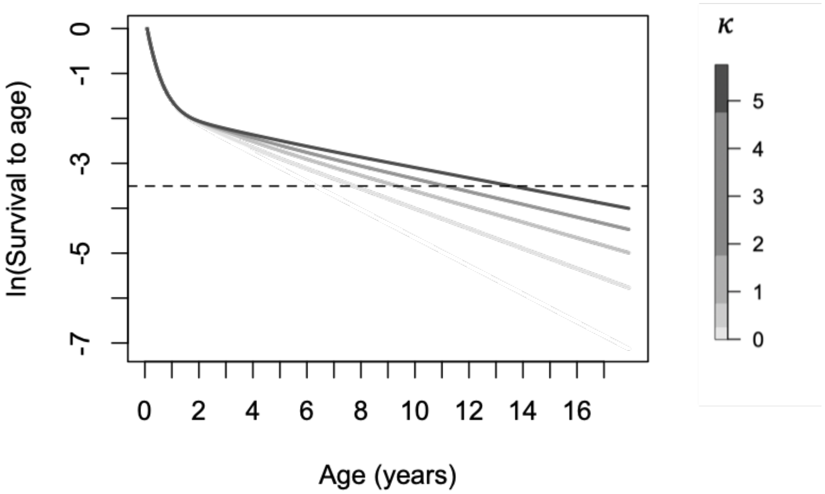


**Supplementary Figure 5.** The emergent pattern of ln(survival) to age at different levels of $\kappa$. Dashed horizonal line shows the cutoff past which less than 3% of individuals will survive. For all results here, $h = 8$ and $\tau = 16.85$°C.

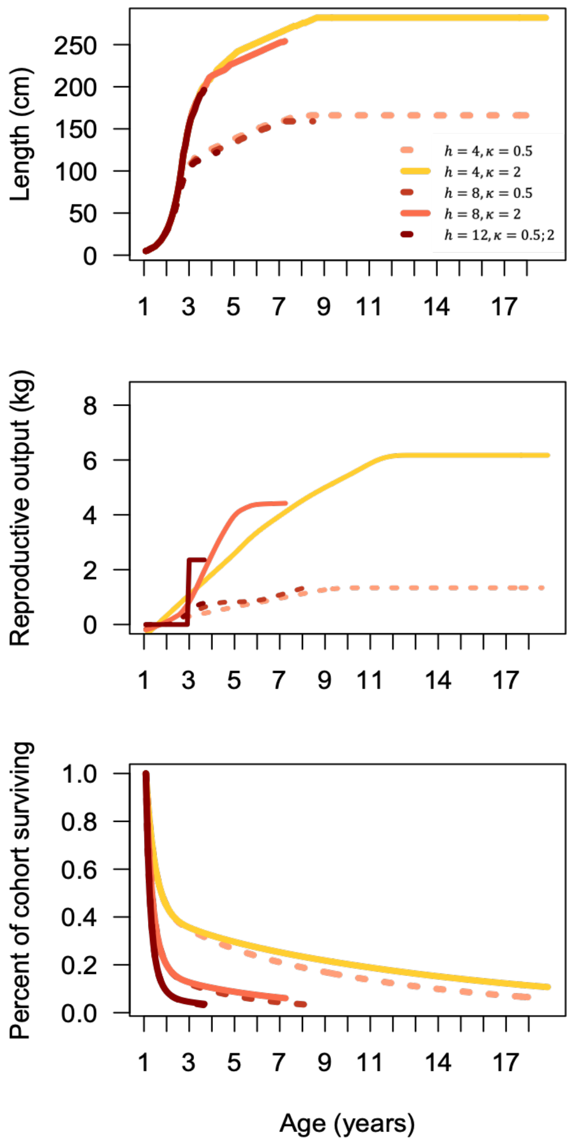


**Supplementary Figure 6.** The emergent patterns of growth, reproduction, and survival that emerge when $h$ varies. We contrast two values of $\kappa$ to highlight that it is still the main driver of variation in body size, although $h$ affects lifespan. In all scenarios, $\tau =$ 16.85℃. Note that for results presented elsewhere, $h$ = 8 except where noted.

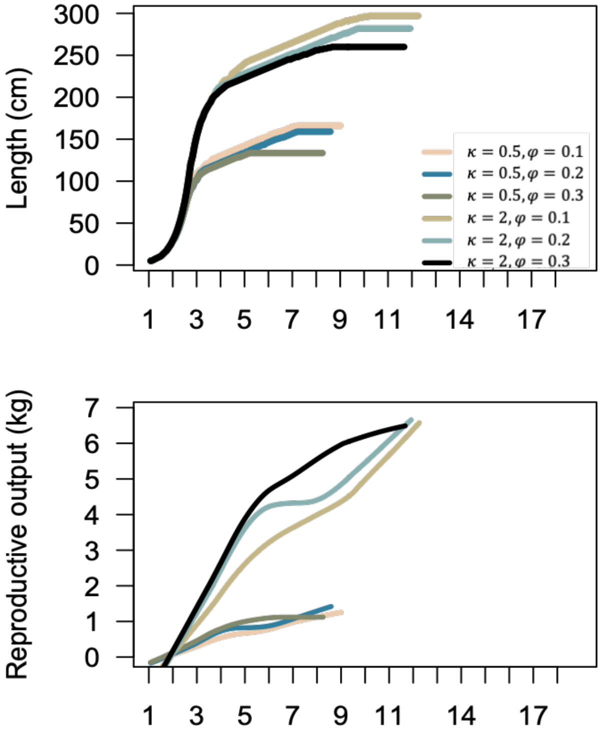


**Supplementary Figure 7.** Growth curves showing the effect of changes in the reproductive capacity $\varphi$, for two values of $\kappa$. As the reproductive constraint increases (the fraction of the body cavity that can be devoted to gonad decreases from 0.2 to 0.1), selection favors the evolution of larger body sizes. In all scenarios, $\tau = 16.85$℃. Note that for all results presented elsewhere, $\varphi = 0.2$.

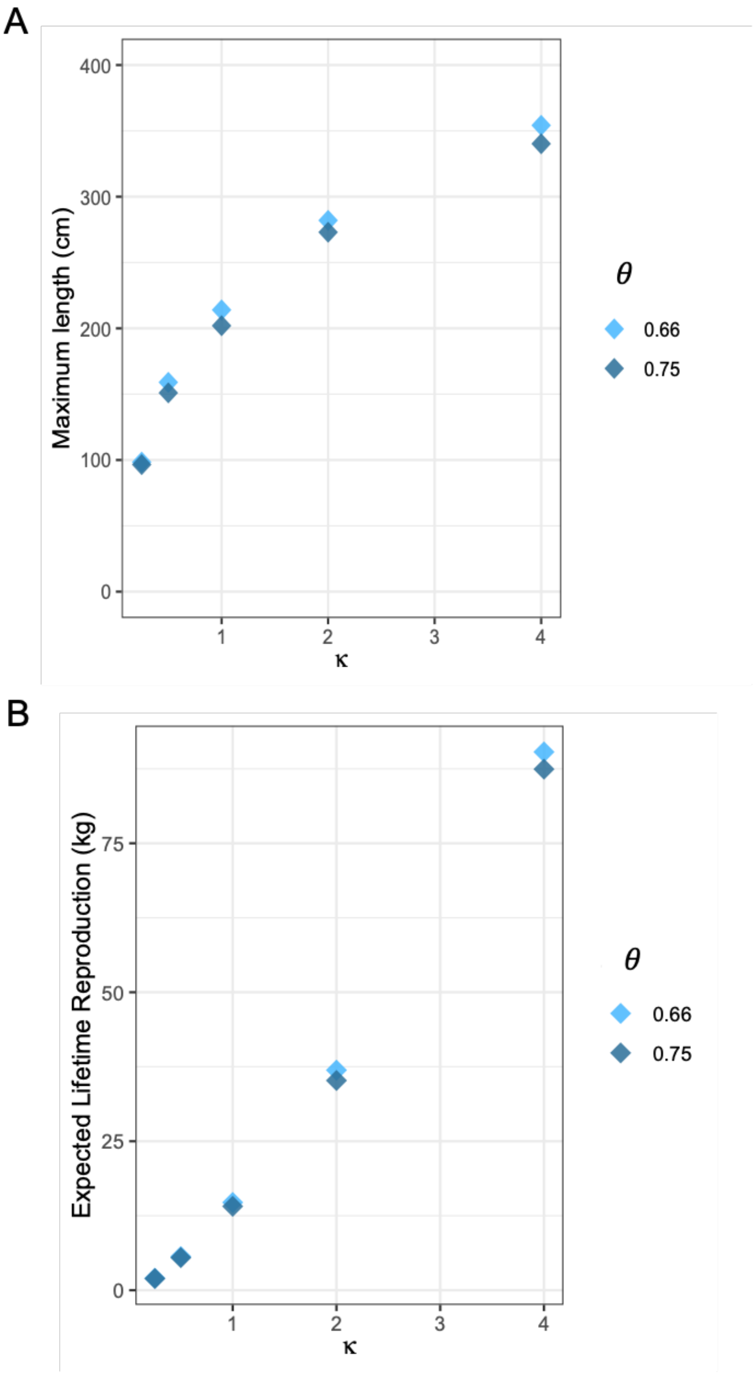


**Supplementary Figure 8.** Body size and lifetime reproduction for several values of $\kappa$ and $\theta$ (the allometric exponent in the metabolic cost function). For all scenarios, temperature is constant year-round at 21.85℃.

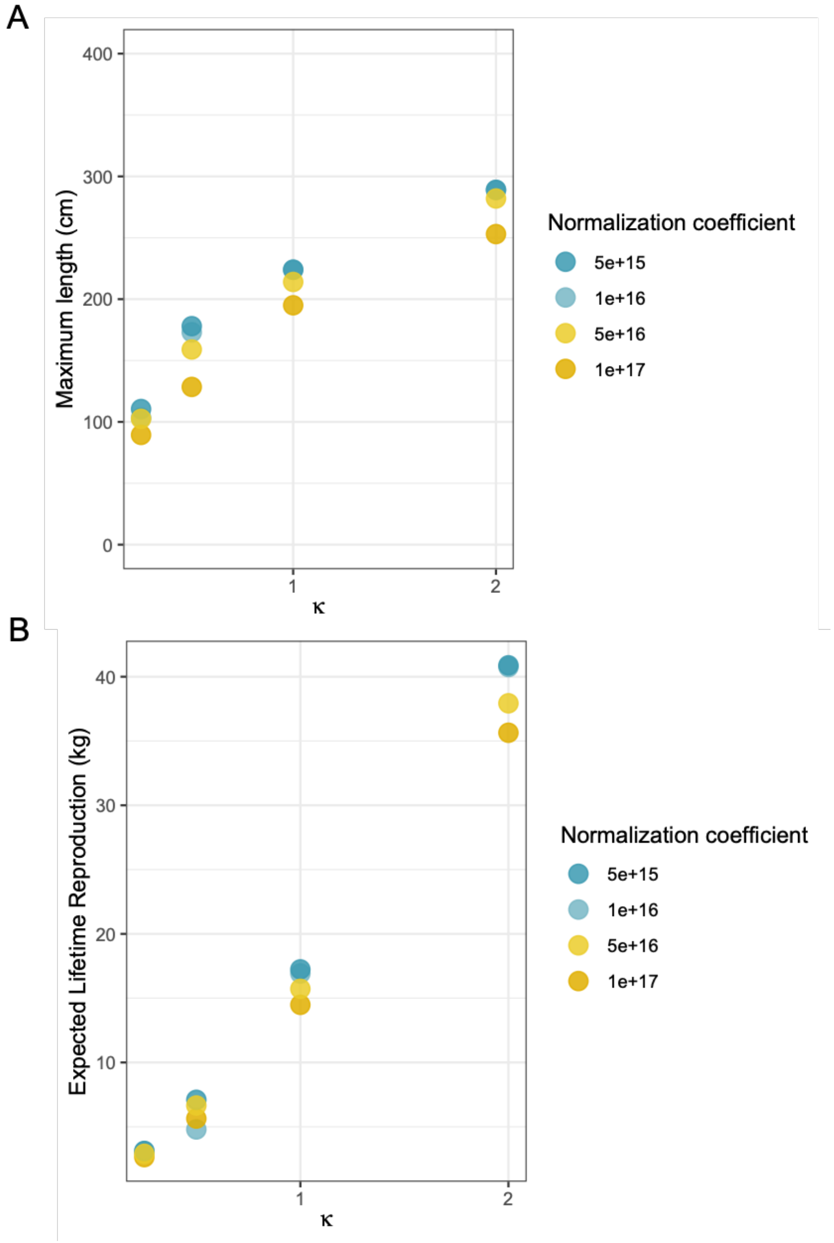


**Supplementary Figure 9.** Body size and lifetime reproduction for several values of $\kappa$ and $c$ (the normalization constant in the metabolic cost function). The estimate of $c$ used elsewhere $(5\times 10^{16})$ comes from metabolic rate data from tunas (Clarke and Johnston 1999). For all scenarios, temperature is constant year-round at 21.85℃.
